# Recombinant production of growth factors for application in cell culture

**DOI:** 10.1101/2022.02.15.480596

**Authors:** Meenakshi Venkatesan, Cameron Semper, Rosa DiLeo, Nathalie Mesa, Peter J Stogios, Alexei Savchenko

## Abstract

Culturing eukaryotic cells has widespread applications in research and industry, including the emerging field of cell-cultured meat production colloquially referred to as “cellular agriculture”. These applications are often restricted by the high cost of growth medium necessary for cell growth. Mitogenic protein growth factors (GFs) are essential components of growth medium and account for upwards of 90% of the total costs. Here, we present a set of expression constructs and a simplified protocol for recombinant production of functionally active GFs, including FGF-2, IGF-1, PDGF-BB and TGF-β1 in *Escherichia coli*. Using this expression system, we produced soluble GFs from species including bovine, chicken, and fish. Bioactivity analysis revealed orthologs with improved performance compared to commercially available alternatives. We estimated that the production cost of GFs using our methodology will significantly reduce the cost of cell culture medium, facilitating low-cost protocols tailored for cultured meat production and tissue engineering.

## Introduction

Cellular agriculture is a rapidly emerging field that seeks to leverage biotechnology to produce agricultural products from cell cultures (Post et al., 2020). Study in cellular agriculture has intensified in recent years due to the proposed environmental and ethical benefits compared to traditional agriculture (Mattick, Landis, Allenby, & Genovese, 2015). The field has spurred the development of a range of products (e.g., milk produced using recombinantly-produced protein (Geistlinger, 2017)); however, the primary focus of research in cellular agriculture relates to the development of cell-cultured meat (CCM). Also referred to as cell-based meat or cultivated meat, these products marry cell culture, tissue engineering, and bioprocessing to develop three-dimensional “slab” meat products independent of animals (save for the initial biopsy used for cell line development or primary cell isolation) (Melzener, Verzijden, Buijs, Post, & Flack, 2021). Direct comparison between CCM and traditionally produced meat products predict that an animal-free approach to meat production can reduce greenhouse gas emissions by up to 95%, conserve land and water usage, attenuate the overuse of antibiotics in agricultural settings, and reduce the incidence of zoonotic infections (Godfray et al., 2018). Furthermore, the development of CCM as a *bona fide* food source has the potential to decentralize supply chains and address issues related to food insecurity and sustainability. The potential benefits of CCM have been well studied; however, for these advantages to be actualized there are several technological challenges that continue to remain as significant hurdles.

The core technology at the foundation of CCM production is eukaryotic cell culture. Cell lines or primary cells of bovine, avian, and *teleost* origin are used to produce cell-cultured beef, chicken, and fish, respectively. Within the field of cellular agriculture, and eukaryotic cell culture more broadly, cells are commonly supported with the addition of 5-20% fetal bovine serum (FBS). FBS is obtained from blood and used as a growth supplement to sustain *in vitro* cell culture by providing hormones, lipids, micronutrients, and, most critically, mitogenic growth factors (GFs) that promote cell division (van der Valk et al., 2018). The use of FBS is near-ubiquitous in research settings; however, its continued use within cellular agriculture is seen as problematic for a number of reasons. FBS is costly, it can suffer from batch-to-batch variation that can result in issues of reproducibility or reduced yield, and the fact that it is sourced from fetal calves undermines the ethical advantage afforded by CCM compared to meat produced using traditional agriculture.

Serum-free alternatives to FBS have been commercially available for decades and in the case of chemically defined serum-free media (SFM), they offer improved consistency more compatible with the controlled manufacturing processes adopted in CCM production. Commercially available serum-free options include Essential 8 (Chen et al., 2011), TeSR, and FBM and recent efforts have endeavoured to use these as templates for creating low-cost serum-free formulations tailored for iPSC cells (Kuo et al., 2020) and bovine satellite cells (Stout, Mirliani, White, Yuen, & Kaplan, 2021). While SFM removes the ethical concerns related with the use of FBS and offers improved consistency in cell culture performance, it is even more cost prohibitive than FBS-containing media. At current prices, the use of SFM for the production of CCM creates a significant barrier for competitive entry into consumer markets.

Mitogenic GFs, such as basic fibroblast growth factor (FGF-2) and transforming growth factor β1 (TGF-β1), are the main cost drivers of SFM, estimated to contribute more than 95% of the total cost (Specht, 2020). These signalling proteins bind to cell surface receptors activating several downstream pathways to result in cell proliferation, cell migration, differentiation, and apoptosis, and are added as a supplement to SFM to mimic the GF composition provided by traditional serum additives. While a wide array of GFs are used to support eukaryotic cell culture, many of which are tissue/cell-type specific, there are a subset which are considered essential for supporting cell proliferation in general (Yao & Asayama, 2017). These GFs are key components of SFM used to support cell growth for academic and cellular agriculture applications.

Recombinant production of these GFs is challenging and is a major contributor to their high cost. They are of eukaryotic origin and typically require some level of post-translational modification, such as oxidation or glycosylation, and proper disulfide bond formation is essential for protein folding and maturation into their biologically active conformation. Consequently, production of these proteins in bacterial expression systems has been challenging. Strategies to address this issue include refolding GFs expressed in bacteria from inclusion bodies or using mammalian cell culture systems (e.g., CHO, HEK293) for their production (Alexander, Hesson, Mannarino, Cable, & Dalie, 1992; Zou & Sun, 2004). Both strategies result in high costs and render SFM cost-prohibitive for use in the production of CCM (Tripathi & Shrivastava, 2019).

*Escherichia coli* remains a staple of recombinant protein production and when compatible with a target protein, offers advantages in terms of speed, tractability, and cost (Rosano & Ceccarelli, 2014). Several advancements in strain engineering have made *E. coli* an improved host for production of disulfide-containing proteins. Mutation of *trxB* and *gor* genes (Bessette, Aslund, Beckwith, & Georgiou, 1999), as well as cytoplasmic overexpression of disulfide bond isomerases (e.g., DsbC) (Lobstein et al., 2012) are strategies that have facilitated the expression of disulfide-containing proteins in *E. coli*, culminating in the development of commercial strains such as Rosetta Gami B (Novagen) and SHuffle T7 (NEB), respectively. Additionally, the use of redox fusion partners such as thioredoxin (Trx) (LaVallie et al., 1993), glutathione S-transferase (GST) (Schafer, Seip, Maertens, Block, & Kubicek, 2015), and disulfide bond isomerases (DsbA, DsbC) (Nozach et al., 2013) have greatly improved soluble expression of proteins that contain disulfide bonds in *E. coli*. Despite these advancements, the use of *E. coli* to produce soluble mitogenic GFs remains underexplored as commercial production still relies on refolding in instances where *E. coli* is the chosen expression host.

To facilitate cost reductions in SFM and to advance its adoption in the field of cellular agriculture, we screened a myriad of GF genes, fusion partner combinations and *E. coli* strains to develop a collection of expression systems that allow for production of biologically active mitogenic GFs in *E. coli*. As a result, we were able to identify expression conditions to successfully produce several GFs from a variety of species including those useful in CCM production (bovine, avian, fish). Expressed GFs can be purified using a one-step immobilized metal affinity chromatography (IMAC) purification procedure with an optional additional “clean-up” step. This methodology is feasible for academic labs as well as CCM start-up ventures and results in biologically active GFs able to promote cellular proliferation. Using this methodology, we identified GF orthologs that provide higher potency than commercially available counterparts. Finally, we provide a techno-economic assessment to show that our methodology facilitates significant cost reductions associated with the recombinant production of mitogenic GFs. These results have the potential to significantly reduce the cost of SFM formulations in the field of cellular agriculture and eukaryotic cell culture applications more broadly.

## Results

### Selection of growth factors and expression system components

GFs to target for cost effective production were selected based on their prevalence in FBS or in current SFM-based formulations, and their likelihood of being widely adopted in CCM production. Based on these criteria, we focused on four major GF families represented by fibroblast growth factor (FGF-1 and FGF-2), platelet-derived growth factor (PDGF-BB), insulin-like growth factor (IGF-1 and IGF-2) and transforming growth factor beta 1 (TGF-β1). For each of these families we selected multiple representatives from diverse species within available genome sequences to sample sequence diversity and to expand the repertoire of GFs able to support the growth of cells relevant for cultured meat production. Furthermore, we tested our most successful expression strategies on additional GFs including leukemia inhibitory factor (LIF), epidermal growth factor (EGF) and myostatin/growth differentiation factor 8 (MSTN/GDF8).

We chose to express the GFs in *E. coli*, which remains the most widely used, versatile, and inexpensive recombinant expression system. Our selection of specific *E. coli* strains and expression vectors was driven by the general goal of maximising the production of GFs in a soluble and biologically active form that would also allow for simplified purification using a standard IMAC protocol. Considering the prevalence of intramolecular disulfide bonds in many GF proteins, we selected commercially available *E. coli* strains altered to accommodate the expression of such proteins including Rosetta Gami 2(DE3) (Novagen), Origami B(DE3) (Novagen) and SHuffle T7 express (NEB) in addition to BL21-Gold (DE3) as the hosts to test for recombinant GF production.

For our first attempt, each of the selected GFs was expressed along with an N-terminal polyhistidine (6xHis) tag or the classic redox fusion partner, thioredoxin A (TrxA). To assess the success of these protein production strategies we performed small-scale expression tests of proteins for each family of GF in the selected *E. coli* host strains and screened for the presence of soluble intracellular protein expression using SDS-PAGE (Figure 1A). We achieved limited success with these two fusion partners for some GF families, so we expanded our screen to include additional redox and non-redox fusion partner sequences. For non-redox fusion partners, we used the 97 amino acid SUMO ubiquitin-like protein from *Sacchromyces cerevisiae* (Addgene plasmid #29711) and the B1 domain of the *Streptococcal* protein G (GB1) (Cheng & Patel, 2004), the latter of which has been successfully used to facilitate soluble expression of human EGF in *E. coli* (Zheng, 2016).

**Figure 1.**
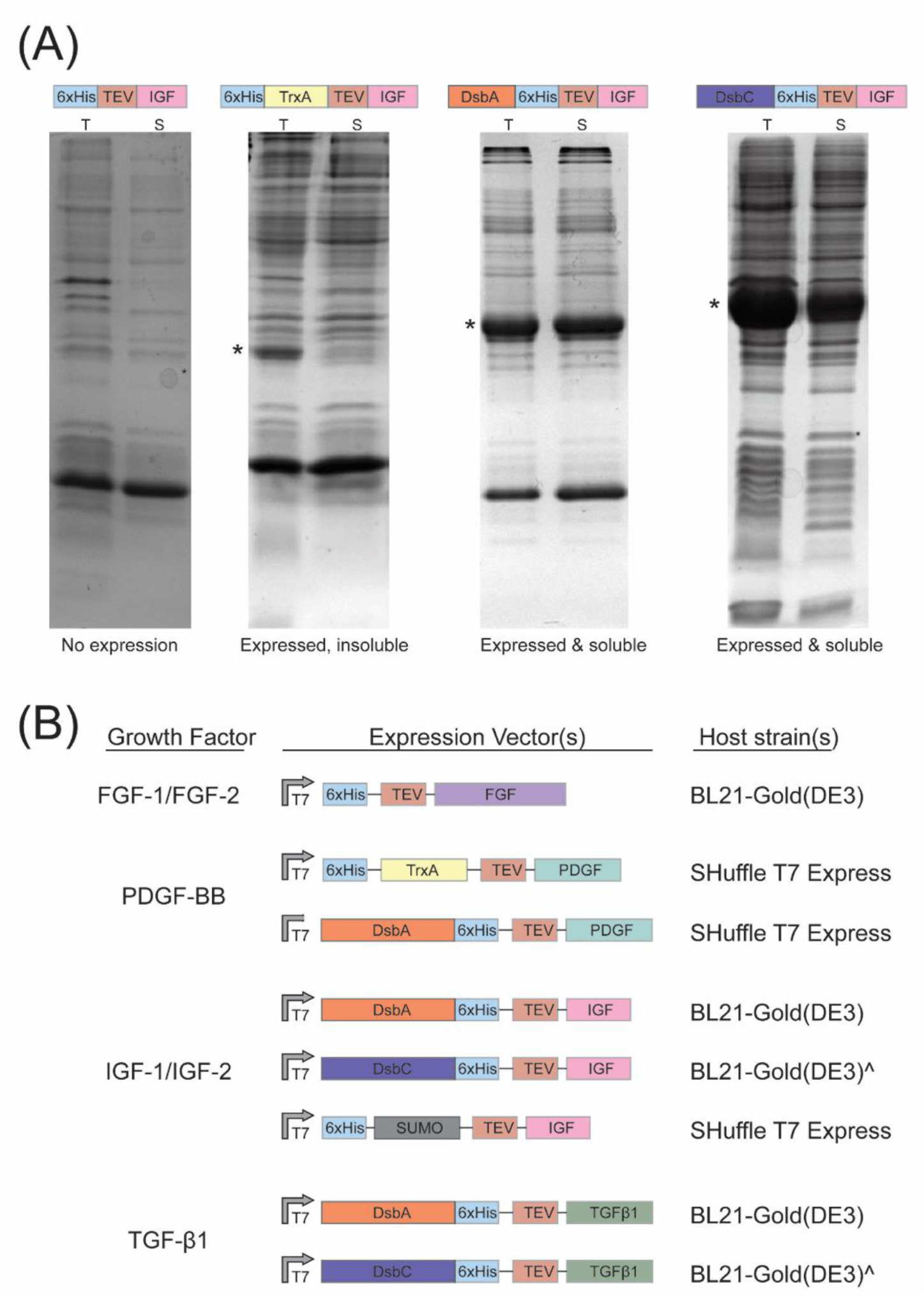
Expression systems for recombinant GF production. (A) Small-scale protein expression screening used to identify the expression vector and host strain combination capable of facilitating cytoplasmic soluble protein expression. The band corresponding to the protein of interest is marked with (*). T - total cell lysate; S - soluble fraction. (B) Expression vector and host strain combinations for successful expression and purification of soluble, bioactive growth factors. (^) denotes instances where the use of SHuffle T7 Express was required for soluble expression of some orthologs.

For additional redox fusion partners, we incorporated disulfide bond isomerases (*E. coli* DsbA or DsbC) that promote formation of disulfide bonds and have been previously described as potent facilitators of soluble expression of disulfide-bond rich proteins in *E. coli*. For this, we modified a previously constructed T7-based expression vector, pMCSG53 (Biancucci et al., 2017; Eschenfeldt et al., 2013), to include either the leaderless DsbA or DsbC protein as an N-terminal fusion partner for the GF protein. These new vectors, pMCSG53-DsbA and pMCSG53- DsbC, are LIC and Gibson Assembly compatible, allowing for high-throughput entry of GF sequences to promote investigation of amino acid sequence diversity within these protein families. All expression constructs included a Tobacco Etch Virus (TEV) protease cleavage site (ENLYFQG) between the tag and GF sequence to allow for removal of the fusion partner after purification.

The expression level of GFs was tested on a small scale (see Material and Methods for details) to identify the most promising strain and construct combinations for soluble recombinant GF expression (Figure 1A). An overview of the most successful expression combination(s) for each GF is summarized in Figure 1B. Below we discuss the outcome of these strategies for each of the selected growth factor families in detail.

### Expression and purification strategy for fibroblast growth factor (FGF-1 and FGF-2)

FGF-1 (acidic FGF) and FGF-2 (basic FGF) are 155 amino acid proteins that exert mitogenic and angiogenic activities on target cells through interaction with fibroblast growth factor receptor (FGFR) and subsequent activation of the downstream RAS-MAPK, P13K-AKT, STAT, and PLCγ signalling pathways (Brewer, Mazot, & Soriano, 2016). FGF-2 also plays a role in tissue development and repair by binding to heparin sulphate cofactor (Koledova et al., 2019). FGF is a critical component of cell culture medium, particularly for maintaining pluripotency of stem cells and their proliferative capacity (Mossahebi-Mohammadi, Quan, Zhang, & Li, 2020). It is also a major mitogen in FBS, required for proliferative activity of cells and is typically present in the range of 10 to 100 ng/mL in cell culture medium (Yang & Xiong, 2012).

Both FGF-1 and FGF-2 have been shown to be prone to proteolytic degradation and denaturation in cell culture, contributing to their relatively short half-life and present a major cost driver for cell culture medium. These undesirable properties can be mitigated through addition of heparin to the growth medium; however, this strategy adds to the overall medium costs and may not meet food safety requirements for CCM applications. FGF variants offering improved thermostability have been identified through protein engineering efforts (Benington, Rajan, Locher, & Lim, 2020; Zakrzewska, Krowarsch, Wiedlocha, & Otlewski, 2004), but their approval for application in CCM products may be complicated due to their “genetically-modified” categorization. With these shortcomings of currently available FGF in mind, we developed a workflow to allow for low-cost bacterial expression and purification of an array of FGF orthologs.

FGF-1 and FGF-2 have been previously produced recombinantly in *E. coli* with a variety of fusion partners (e.g., TrxA, 6HFh8, (Gasparian et al., 2009; Kim et al., 2021)); however, these approaches often require multiple purification steps and the use of specialized chromatography equipment. In contrast, we tested the production of these GFs by a single step IMAC protocol, followed by a TEV protease tag cleavage and removal using the same affinity resin used for initial purification as shown in Figure 2a (see details in Materials and Methods). Additional modifications in our methodology from previous expression strategies for FGF proteins include the use of the synthetic codon-optimized genes and sampling of 21 sequence orthologs (5 FGF-1, 16 FGF-2) to capture a representative level of amino acid sequence diversity within this protein family. Our strategy allowed for successful expression and purification of 5 FGF-1 and 16 FGF-2 from terrestrial (bovine, chicken, sheep, human) and aquatic (Atlantic salmon, Nile tilapia, Chinook salmon) species (Tables S1 & S2). The protein yields of purified FGF-2 ranged from 7 mg/L for *Bos taurus* (bovine) to 36 mg/L for *Oryzias latipes* (rice fish), highlighting significant variation in FGF ortholog yield while expressed under the same general conditions (Figure S1).

**Figure 2.**
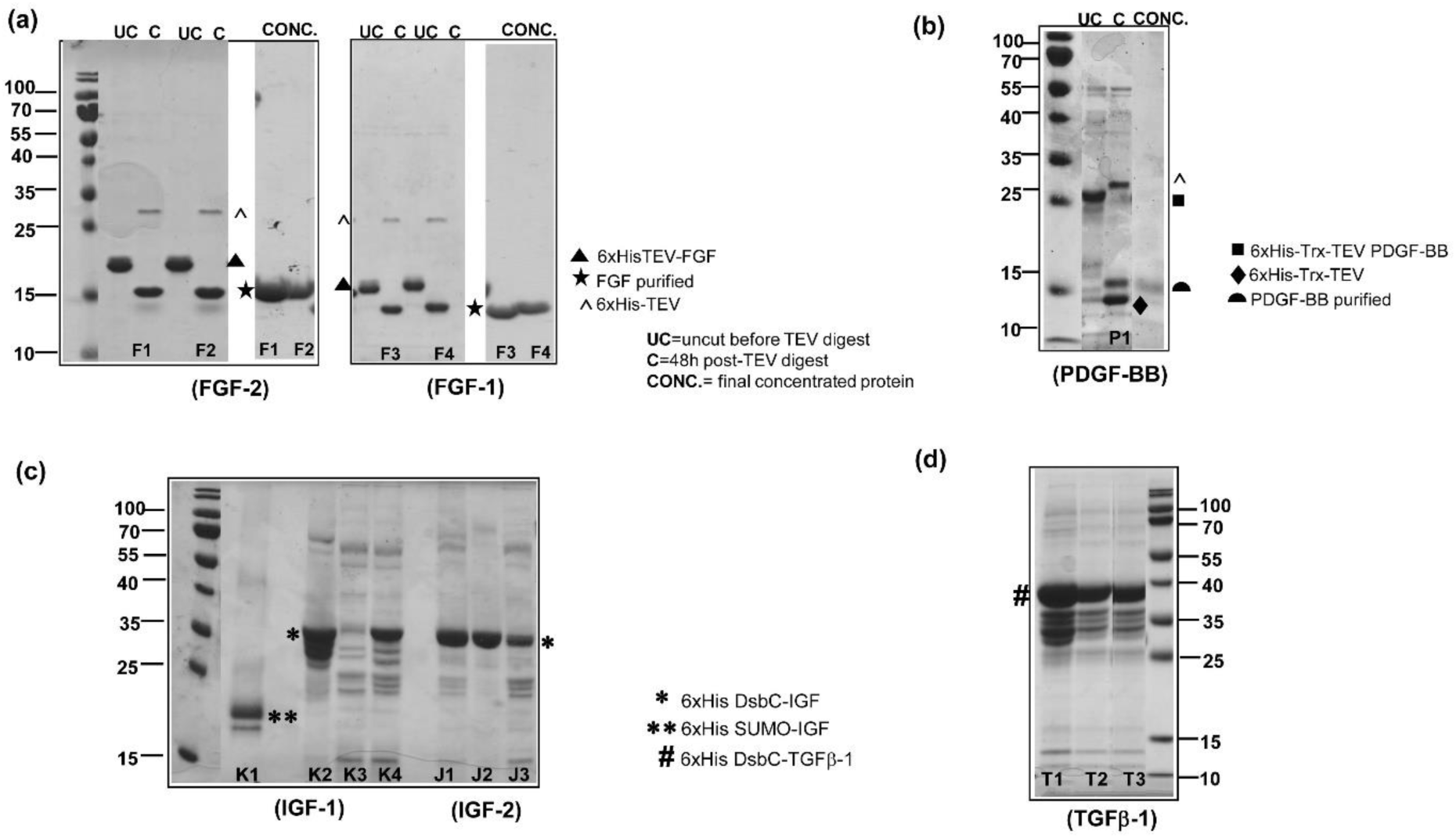
Recombinant GF production. Scale up of protein expressions for **(A)** FGF-2 AND FGF-1 cloned in pMCSG53 vector with N-terminal His6x tag and expressed in BL21(DE3) Gold cells. Targets include **F1** (FGF2_Atlantic salmon); **F2** (FGF2_Pufferfish); **F3** (FGF1_Sheep); **F4** (FGF1_Bovine) **(B)** PDGF-BB expressed in SHuffle T7 express cells. Target shown is **P1** (PDGFBB_Cormorant) **(C)** IGF-1/IGF-2 cloned in pMCSG53-His6x-DsbC /pMCSG53-His6x-SUMO and expressed in SHuffle T7 express cells. Targets include (**K1**) IGF1_Bovine (SUMO-His6x tag); (**K2**) IGF1_Bovine (DsbC-His6x tag); **(K3**) IGF1_Goose; **(K4**) IGF1_Frog; (**J1**) IGF2_Human; (**J2**) IGF2_Bovine; (**J3**) IGF2_Nile tilapia **(D)** TGFβ-1 cloned in pMCSG53-His6x-DsbC and expressed in SHuffle T7 express cells. Targets shown are TGFβ1_human (**T1**); TGFβ−1_bovine (**T2**); TGFβ−1_chicken (**T3**); TGFβ−1_little egret (**T4**). **UC***=uncut before TEV digest;* **C***=48h post-TEV digest;* TEV protease runs at 25 kDa (marked with ^). After the TEV digest and a second Ni-NTA affinity chromatography step, the concentrated, purified FGF-2/FGF-1 runs at 15 kDa on an SDS-PAGE (marked with ) shown in **(A)**. PDGF-BB runs at 15 kDa corresponding to the monomer (marked with ⊇) shown in **(B)**. DsbC fusion IGF-1/IGF-2 runs at 35 kDa (marked with *). IGF1-SUMO runs at 20 kDa (marked with **), as seen in **(C)**. DsbC-TGFβ-1 runs at 40 kDa (marked with *#*).

### Expression and purification strategy for Platelet-derived growth factor (PDGF-BB)

Platelet-derived growth factors (PDGF) are potent mitogens for a variety of cells including fibroblasts, smooth muscle cells, connective tissue cells, and bone and cartilage cells. PDGF has been shown to increase proliferation of mouse myoblasts and is a key component for the production of CCM (Albrecht & Tidball, 1997). Isoforms (A, B, C, and D) of PDGF form homo- or heterodimers stabilised by intermolecular disulfide bonding. All four PDGF isoforms feature a highly conserved 100 amino acid GF domain (Breitkopf et al., 2005). Dimerized PDGF binds to two cognate tyrosine kinase receptors, platelet-derived growth factor receptor α (PDGFRα) and PDGFRβ, initiating intracellular signal transduction pathways. Examples of pathways involved in response to PDGF-BB activation include MEK/ERK, Src, and PI3K/AKT (Fredriksson, Li, & Eriksson, 2004).

Commercially available human PDGF-BB is produced recombinantly in *E. coli;* however, previously published protocols are plagued by low yields, or rely on purification from inclusion bodies under denaturing conditions with the requirement of subsequent refolding to obtain bioactive PDGF-BB (Alexander et al., 1992). To improve upon existing PDGF expression procedures we tested expression of 25 PDGF-BB orthologs (avian, mammalian, reptile, and aquatic) in fusion with N-terminal 6xHis tag and 6xHis-TrxA or 6xHis-DsbA double tags. The 6xHis-TEV-PDGF-BB fusions were individually expressed in BL21(DE3)-Gold, Rosetta Gami B, and SHuffle T7 *E.coli* host strains, but only 1 out of 25 GFs showed soluble expression under these conditions. In contrast, the use of an N-terminal 6xHis-TrxA double fusion resulted in 21 out of 25 PDGF-BB produced in a cytoplasmic soluble form (Table S2). We scaled up protein expression and purified 12 of these PDGF-BB orthologues ranging from bovine, bony-tongue fish, cormorant, eagle, cobra, green anole and sea turtle and achieving protein yields ranging from 2-5 mg/L (Figure S1). The 6xHis-TrxA tag was efficiently cleaved from PDGF-BB using TEV protease (Figure 2B); however, in a subset of the orthologs that were purified we observed co-elution of TrxA and PDGF-BB after the second Ni-NTA purification step. Adding a size-exclusion chromatography step was successful in separation of PDGF-BB from TrxA in cases where this tag remained associated with PDGF-BB after TEV protease digestion (Figure S3).

The four PDGF-BB orthologues that were not produced as soluble proteins using the pET-TrxA fusion were all sourced from *Teleost* (ray-finned fish) species. Fusion of these orthologues to an N-terminal DsbA tag allowed for their soluble expression and purification. Using this approach, the PDGF-BB from climbing perch (*Anabarillus testudineus*) was purified at a yield of 7 mg/L. Analysis of all the recombinantly purified PDGF-BB proteins via non-reducing SDS-PAGE confirmed the expected pattern of protein dimerization (Figure S3).

### Expression and purification strategy for Insulin-like growth factor (IGF-1 and IGF-2)

IGF-1 and IGF-2 are 70 amino acid disulfide-rich proteins with a similar structure to the peptide hormone insulin (Brown et al., 2008; Vajdos et al., 2001). IGF signalling is central to pathways that control cell growth, maturation, and proliferation. IGF-1 is a major component of FBS and is widely used in cell culture media including in Essential 8 (E8) medium as well as in many different formulations used to promote myogenesis and the development of skeletal muscle (Hakuno & Takahashi, 2018; McCubrey et al., 1991; Yu et al., 2015). IGF-1 and IGF-2 function on cells by binding to receptor-tyrosine kinases (RTKs), the IGF-1 receptor (IGF1R) and IGF-2 receptor (IGF2R), respectively. IGF-1 and IGF-2 can also bind the insulin receptor, though they do so with lower affinity compared to their interactions with their cognate receptors. IGF-1 and IGF-2 initiate signal transduction through the PI3K/AKT and ERK1/2 signalling pathways resulting in stimulation of cell growth and proliferation (Peruzzi et al., 1999). Commercially available IGF-1 is typically produced in *E. coli* as inclusion bodies, purified under denaturing conditions and then refolded into the bioactive form. Recent studies have reported that DsbA (Emamipour, Vossoughi, Mahboudi, Golkar, & Fard-Esfahani, 2019) or 6HFh8 (a six histidine tag followed by the Fh8 peptide from *Fasciola hepatica*) (Kim et al., 2021) fusion partners promote the soluble expression of mammalian IGF-1 in *E. coli*.

Based on our success using the TrxA fusion tag to facilitate PDGF-BB expression, we tested the same approach for expression of 15 IGF-1 and IGF-2 orthologs derived from mammalian, avian and fish species. This strategy did not result in soluble expression of IGF based on small-scale expression testing (Figure 1A). Next, we assessed the ability of DsbC, DsbA, and SUMO fusion tags to promote soluble expression of IGF in *E. coli*. Expression of IGF as a DsbC-IGF fusion resulted in significant increases in the production of soluble GF protein (Figures 1A, 2C). With this expression system, all the selected IGF orthologs (human, bovine, fish) were produced primarily as soluble fusion protein (6 IGF-1 and 3 IGF-2), (Figure S1). Yields of DsbC-IGF-1 fusion protein ranged from 6 to 10 mg/L. The DsbA-IGF-2 fusion had similar results in terms of protein solubility, but the overall expression level and yield was generally less than compared to expression of DsbC-IGF-1 fusions. The expression of SUMO-IGF-1 fusions also resulted in production of soluble IGF protein but only in the SHuffle T7 *E. coli* host strain. TEV protease cleavage of the DsbA/DsbC fusion tags was efficient; however, similar to what we observed for some TrxA-PDGF-BB fusion proteins, the cleaved fusion partner co-eluted with IGF-1 after the second Ni-NTA chromatography step. Anion exchange chromatography (MonoQ) resulted in some degree of separation, but the addition of this step may be problematic at production scales necessary for cellular agriculture applications. To determine if separation of the fusion DsbA/DsbC tag from IGF-1/IGF-2 was a necessary step for bioactivity, we assayed the ability of these fusion GFs to support cell growth and to ascertain if the presence of an N-terminal DsbA/DsbC/SUMO tag had an impact on their functional bioactivity (discussed in sections below).

### Expression and purification strategy for Transforming growth factor beta 1 (TGF-β1)

TGF-β1 belongs to the transforming growth factor superfamily which consists of isoforms TGF-β1-3 as well as of other signalling proteins such as myostatin, growth differentiation factor (MSTN or GDF8), and bone morphogenetic factor (BMP) (Poniatowski, Wojdasiewicz, Gasik, & Szukiewicz, 2015). TGF-β GF are synthesized as a larger precursor protein and then proteolytically cleaved into a 112 amino acid active GF (Derynck et al., 1985). They are rich in cysteine residues and contain multiple intramolecular disulfide bonds participating in formation of a cysteine knot (Hinck et al., 1996). Additionally, disulfide bonding results in the formation of a covalently-linked homodimer that is critical for biological activity. TGF-β1 is a key component of the growth medium used for skeletal muscle growth and differentiation and is critical for maintaining the pluripotency of stem cells (Mullen & Wrana, 2017; Scharenberg et al., 2014). For these reasons, it is a major component of growth medium optimized for production of CCM (Boudreault, Tremblay, Pepin, & Garnier, 2001).

Recombinant production of TGF-β1 has traditionally relied on expression using eukaryotic expression systems such as Chinese hamster ovary (CHO) or human embryonic kidney (HEK293) cells, contributing to its high cost and role as a major cost driver in cell culture media (Zou & Sun, 2004). A recent report detailed successful soluble expression of TGF-β3 in *E. coli*in fusion with a TrxA tag (Kuo et al., 2020), but to our knowledge, soluble expression of TGF-β1 in *E. coli* has not been reported. Based on our success in using the TrxA fusion partner to facilitate soluble expression of PDGF-BB, and reports detailing soluble expression of TGF-β3 using this fusion partner, we attempted to express 10 TGF-β1 orthologs (human, bovine, fish, avian) with an N-terminal TrxA fusion. While we did observe expression of a protein species of appropriate size, none of the TrxA-TGF-β1 orthologs were expressed as a soluble protein. Next, we tested GB1 tag as the TGF-β1 fusion partner, but the results were similar with the expressed protein found only in the insoluble fraction. In contrast, expression of TGF-β1 as a DsbA or DsbC fusion protein resulted in several TGF-β1 orthologs (human, bovine, fish, avian) expressed in cytoplasmic soluble fraction. Yields of purified DsbC-TGF-β1 fusion protein ranged from 4.5 to 6 mg/L (Figure 2d). As observed with the IGF-1/IGF-2 GFs, separation of DsbC/DsbA from TGF-β1 after TEV-digest required an anion exchange chromatography step.

### Expression and purification of additional growth factor families

Our success with recombinant expression of GF that are the key components of SFM prompted us to try similar strategies for expression of additional GFs that have utility in CCM production. Epidermal growth factor (EGF) is used in differentiation of satellite cells into myotubes and supports the proliferative capacity of mammalian adipose-derived stem cells (Ai et al., 2017). A previous study showed that a GB1 fusion supported soluble expression of human EGF in *E. coli*(Zheng, 2016). We tested this system on EGF orthologs from fish species and confirmed its ability to promote soluble expression of EGF from *Teleost* fish species (Supplementary figure 5). We were able to produce soluble GB1-EGF from *Betta splendens* with a yield of 13.5 mg/L.

Leukemia inhibitory factor (LIF) is used in supporting cell culture of bovine satellite cells (Spangenburg & Booth, 2002). We had success in achieving soluble expression of LIF orthologs from human, bovine, fish, and avian species by using an N-terminal 6xHis-GB1 double fusion partner (Figure S3, Figure S5). Myostatin (GDF8) is a member of the TGF-β superfamily and is important in regulating muscle development and skeletal muscle mass. Similar to our results with TGF-β1, an N-terminal DsbC fusion facilitated soluble cytoplasmic expression of myostatin (GDF8) orthologues from human and bovine species (Figure S5.). These results further validate our developed methodology for GFs and its application to several GF families relevant to culture media formulation.

### Validating the functional bioactivity of the recombinantly purified growth factors

We tested the purified recombinant GFs for their ability to stimulate the proliferation of NIH-3T3 mouse fibroblast cells in a dose-dependent manner using a colorimetric MTT assay. This assay follows the cellular metabolic activity as a measure of viable proliferating cells. To assess the dose-dependent behaviour, serum-starved fibroblast NIH-3T3 cells were individually treated with varying concentrations of each purified GF. All of the recombinantly purified GFs tested demonstrated the ability to support similar and, in some cases higher, cell proliferation than their commercially available mammalian counterparts. We report the cell proliferation capacity of cultures containing each recombinantly produced GF as a fold-change when compared with that of the non-supplemented, serum-starved cells (negative control) under similar experimental conditions (Figure 3). Cells grown on culture medium supplemented with 10% commercial FBS served as the positive control.

**Figure 3.**
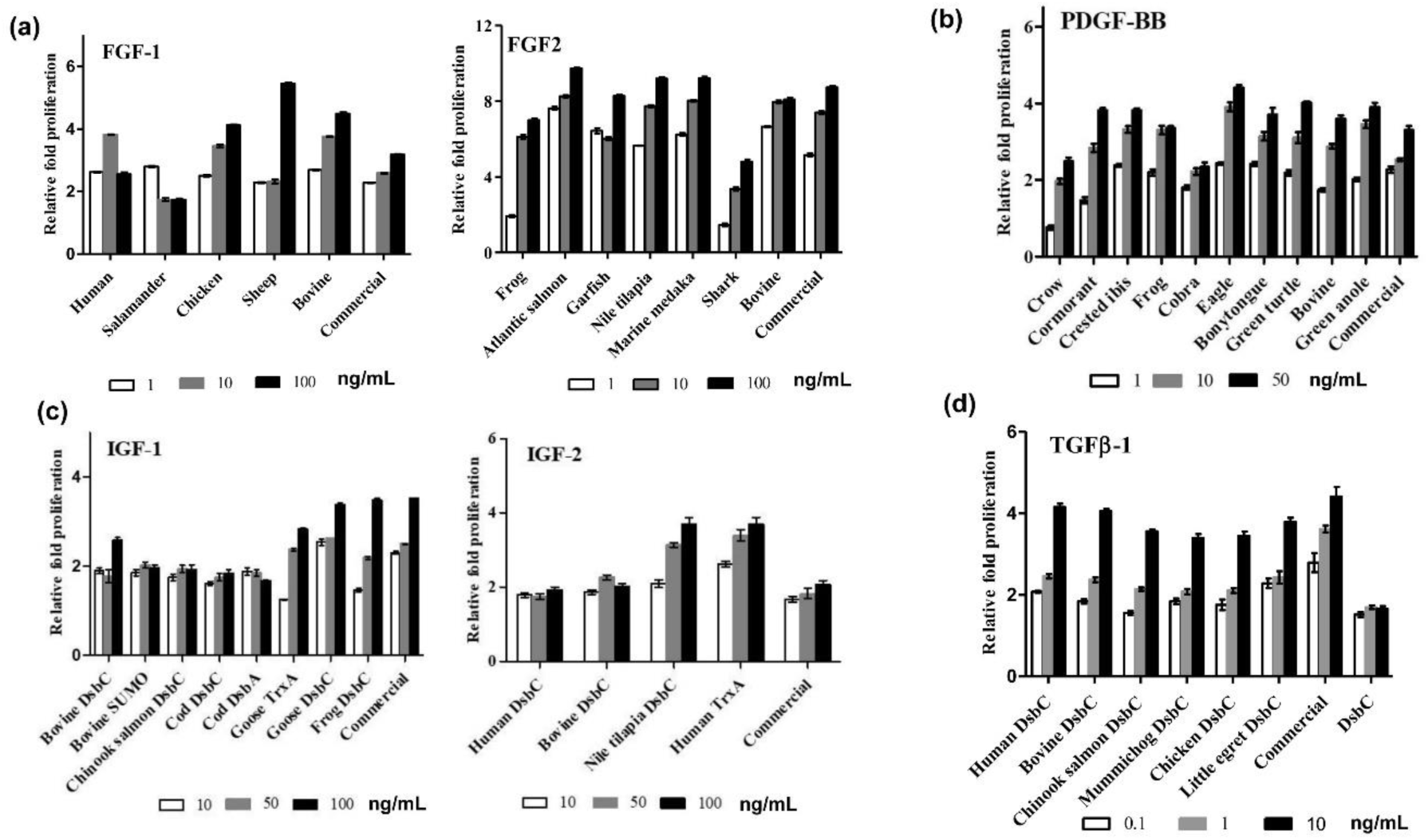
Functional bioactivity validation by colorimetric MTT assay. For each growth factor target, cell proliferation is plotted as relative fold-change when compared with that of the untreated serum-starved cells under similar experimental conditions. Absorbance readings at 540 nm were recorded in six replicates and averaged. Data plotted as mean ± standard deviation (SD) using Graphpad Prism v5.0. Error bars represent the SD calculated for the replicates. Growth factor concentrations are in ng/mL.

The culture medium treated with FGF-2 orthologs derived from bovine (*Bos taurus*), Atlantic salmon (*Salmo salar)*, Marine medaka (*Oryzias melastigma*), and Nile tilapia *(Oreochromis niloticus*), at 10 ng/mL, all showed similar-fold proliferation (8.0 to 8.3-fold for the aforementioned FGF-2 orthologs versus 7.4-fold proliferation with commercial human FGF-2). In the case of FGF-2 orthologs derived from bovine and Atlantic salmon, cells treated with these FGF-2 variants showed 1.5x higher proliferation (6.8-fold and 7.6-fold, respectively), at a much lower (1 ng/mL) FGF-2 concentration compared to cells grown in culture medium supplemented with commercial human FGF-2 (5-fold change compared to serum starvation). If this fold-change could be equivalently achieved in bovine myoblast cells or fish cells, it could have a significant impact on the cost incurred for culture media formulation in CCM production. According to our results, the FGF-2 orthologs we tested can accommodate up to a 10X reduction in the amount of FGF-2 added while still promoting similar levels of cell proliferation.

We observed a similar phenomenon when assaying recombinant FGF-1. FGF-1 orthologs derived from human (*Homo sapiens*) and bovine (*Bos taurus*) showed 1.6x higher proliferative capacity when compared to the commercial human FGF-1 at 10 ng/mL. The FGF-1 ortholog derived from chicken (*Gallus gallus*) showed 3.5-fold proliferation at a concentration of 10 ng/mL in comparison to 2.5-fold with commercial human FGF-1. At 100 ng/mL, FGF-1 from sheep achieved 1.8x higher proliferative capacity as compared to commercial human FGF-1.

PDGF-BB orthologs derived from bovine (*Bos taurus*), crested ibis (*Nipponia nippon*), frog (*Xenopus tropicalis*), eagle (*Haliaeetus albicilla*), and green anole (*Anolis carolinensis*) showed comparatively higher fold-proliferation (3.5- to 4.0-fold) at a much lower concentration of 10 ng/mL in comparison to commercial PDGF-BB (3.5-fold) proliferation at 50 ng/mL, which is the typical culture medium concentration for PDGF-BB. The ability to produce these PDGF-BB orthologs at a low cost and their potential to sufficiently stimulate proliferation at lower concentrations makes them key candidates for advancing cost reductions in CCM production.

All the orthologs of IGFs derived from expression with the different fusion partners (SUMO, DsbC, DsbA) showed similar fold-proliferation as the commercial human IGF-1/IGF-2. Recombinantly purified bovine IGF-1 (DsbC fusion) showed 2.6-fold versus 3.5-fold proliferation for commercial human IGF-1 at 100 ng/mL concentration when compared to untreated serum-starved cells. Purified IGF-2 orthologs derived from human and bovine both showed similar 2-fold proliferation as the commercial human IGF-2 at a concentration of 100 ng/mL. Interestingly, IGF-2 ortholog from Nile tilapia (*Oreochromis niloticus*) showed 1.8x higher fold proliferation than the commercial human IGF-2 (3.6-fold and 2-fold respectively at a concentration of 100 ng/mL).

TGF-β1 orthologs from human and bovine both showed 4-fold proliferation as compared to 4.4-fold from commercial human TGF-β1 at a concentration of 10 ng/mL. TGF-β1 from chicken (*Gallus gallus*), northern pike (*Egretta garzetta*), Chinook salmon (*Oncorhynchus* tshawytcha) and mummichog (*Fundulus* heteroclitus) showed 3.5-fold, 3.8-fold, 3.6-fold, and 3.4-fold respectively, proliferation at a concentration of 10 ng/mL when compared with that of untreated cells. Taken together, our results with IGFs and TGF-β1 demonstrate that while it is possible to remove the 6xHis-DsbC, DsbA or SUMO fusion partners, the presence of the fusion tags does not interfere with bioactivity of IGF-1/IGF-2 or TGF-β1 in cell culture medium.

In addition to measuring the ability of our recombinantly produced GFs to stimulate cell proliferation, we conducted Western blot analysis to observe activation of the downstream cell signalling pathways (ERK, MAPK, Akt) that regulate cellular proliferation. Activation of ERK1/2 through phosphorylation is the key determinant of the cellular response to FGF, PDGF-BB, and IGF growth factor signalling (Zhu, Duchesne, Rudland, & Fernig, 2010). NIH-3T3 cells treated with the recombinantly purified FGF-2, PDGF-BB, and DsbC-IGF-1/IGF-2 fusion orthologs were analyzed for the detection of phosphorylated ERK1/2 (p-ERK1/2). ERK1/2 requires a dual phosphorylation of conserved threonine and tyrosine residues to be activated. Our Western blot results showed bands at 42 and 44 kDa, indicative of phosphorylated ERK1/2, thus complementing the results of our cell proliferation assays and validating that our recombinant GFs activate the expected intracellular signalling pathways (Figure 4).

**Figure 4.**
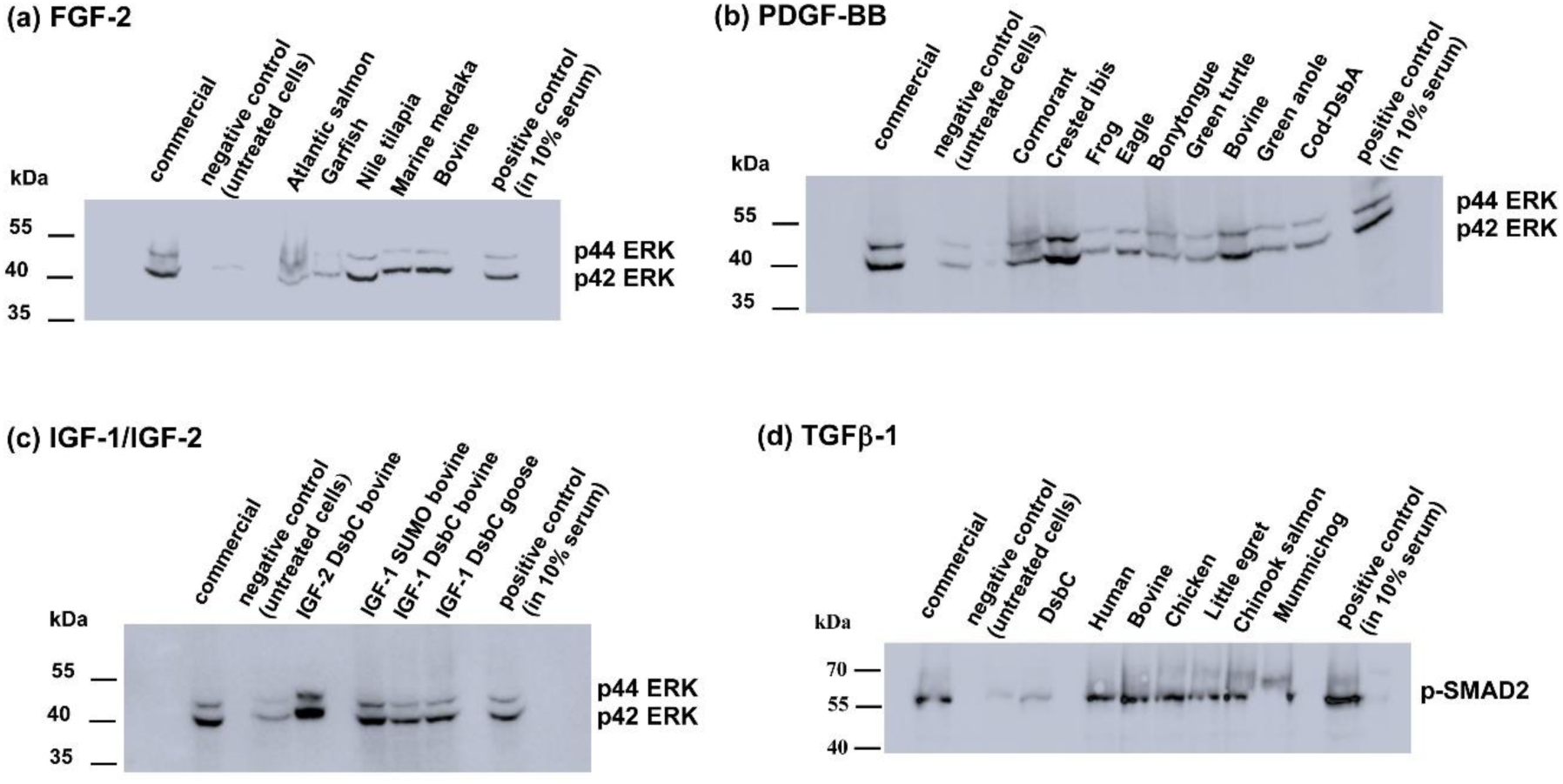
Western blot analysis of cell extracts from NIH-3T3 cells treated GFs. Cells treated with **(A)** FGF-2 targets at 10 ng/mL for 24h at 37 °C **(B)** PDGF-BB targets at 50 ng/mL for 24h at 37 °C **(C)** IGF-1/IGF-2 targets at 100 ng/mL for 24h at 37 °C. All the lysed cell extracts were analyzed for phosphorylated p44/p42 ERK1/2 (p-ERK1/2, 42, 44 kDa). Negative control (untreated cells) shows no detection of phosphorylated ERK1/2, while all the recombinantly purified FGF-2, PDGF-BB, IGF-1, IGF-2 targets and their commercial counterparts show the presence of bands approximately at 42 kDa and 44 kDa **(D)** TGFβ−1 targets at 10 ng/mL for 45 min at 37 °C and analyzed for phosphorylated SMAD2 (p-smad2; 60 kDa). Negative control (untreated cells) shows no detection of p-smad2 band while all the recombinantly purified TGFβ−1 targets and commercial TGFβ−1 show the presence of a band at 60 kDa.

Phosphorylation of SMAD proteins is an event that occurs after TGF-β1 binds to the TGFβ-receptor at the cell surface, thereby promoting cell proliferation (Schmierer & Hill, 2007). To confirm that our DsbC-TGF-β1 fusion proteins were functionally active, we performed Western blotting against phosphorylated SMAD2 (p-SMAD2). Our results showed that the recombinantly purified DsbC-TGF-β1 GF were capable of inducing phosphorylation of SMAD2, validating this GF retains the proper signalling functionality even with the DsbC fusion tag. (Figure 4).

### Techno-economic analysis of “in-house” growth factor production

We sought to develop an economic assessment for production of GFs using the methodology in this study. An estimate based on laboratory consumables costs and labour cost showed that we could achieve recombinant production of growth factors for CAD$ 10.22 per milligram of purified protein (Table 1). A one-time cloning cost incurred for each GF target protein (inclusive of gene synthesis, sequencing primers and PCR) was calculated as approximately CAD$ 60.00. A more rigorous breakdown for the total cost estimation analysis is presented in the Supplementary section (Table S1).

**Table 1.** Production cost of “in house” growth factors: All pricing in CAD and per milligram purified protein. Total production cost of 1225.90 CAD includes all lab consumables and labour costs (**see Supplementary Table 1**) and estimated for 12 litre production scale of E.coli culture growth. Calculations based on total yield of 120 mg purified protein for all except PDGFBB for which the total yield is 60 mg from a 12L culture.

| Growth factors | Protein production costs ("in-house"), (CAD) | Cost "in-house"/mg protein (CAD) | Cost PeproTech /mg protein (CAD) | Cost R&D systems /mg protein (CAD) | Expression system used, commercial suppliers |
| --- | --- | --- | --- | --- | --- |
| TGFβ-1 | \$1225.90 /120 mg protein | \$ 10.22 | \$ 6,240.00 | \$ 7,150.00 | CHO cells, HEK293 cells |
| FGF-2/FGF-1 | \$1225.90/120 mg protein | \$ 10.22 | \$ 960.00 | \$ 2,600.00 | *E. coli* cells |
| IGF-1/IGF-2 | \$1225.90/120 mg protein | \$ 10.22 | \$ 300.00 | \$ 766.00 | *E. coli* (from inclusion bodies) |
| PDGF-BB | \$1225.90 /60 mg protein | \$ 20.43 | \$ 4,680.00 | \$ 5,500.00 | *E. coli* (from inclusion bodies) |

Earlier studies had estimated that FGF-2 and TGF-β1 can account for as much as 95% of the costs of SFM when sourced from traditional commercial suppliers (Specht, 2020). Using our “in-house” methodology, the overall cost contribution of GFs (FGF-2 and TGF-β1) to the total cost of SFM can be substantially reduced (Table 2a). Tables 2b and 2c summarize a price comparison in the contribution of GF costs, specifically shown for Essential 8 medium demonstrating an 11-fold reduced culture medium cost and a significant reduction in the GF cost contribution to the total media cost from 95% to 4%.

**Table 2(a).** Pricing comparison of TGFβ1 and FGF-2 of “in-house” versus commercial (R&D systems commercial source as reference). Below mg/L growth factor calculated based on TGFB1 and FGF-2 composition in Essential 8 media

| Growth factors | Final conc. (in mg/L) in CCM | Cost/L ("in house"), CAD | Cost/L (R&D Systems), CAD | Pricing comparison, fold-change |
| --- | --- | --- | --- | --- |
| TGFβ1 (10 ng/mL) | 0.01 | \$ 0.10 | \$ 72.00 | **720x ↓** |
| FGF2 (100 ng/mL) | 0.1 | \$ 1.02 | \$ 260.00 | **255x ↓** |

**Table 2(b).** Pricing comparison of total cost of cell culture medium (DMEM basal media and growth factor components). Media pricing was estimated per litre of Essential 8 medium composition. DMEM basal media pricing was derived from thermoFisher vendor (cat. no. 11995065) at CAD$ 15.53/500 mL. *The cell culture medium cost per litre is 11-fold cheaper using our “in-house” growth factors*.

| Growth factors | Final conc (mg/L) in cell culture media | Cost/L ("in house"), CAD | Cost/L (R&D Systems), CAD | Pricing compare, fold-change |
| --- | --- | --- | --- | --- |
| TGFβ1 (10 ng/mL) | 0.01 | \$ 0.10 | \$ 72.00 | **720x ↓** |
| FGF2 (100 ng/mL) | 0.1 | \$ 1.02 | \$ 260.00 | **255x ↓** |
| | DMEM basal media | \$ 31.06 | \$ 31.06 | **N/A** |
| | **TOTAL COST** | **\$ 32.18** | **\$ 363.06** | |

**Table 2(c).**
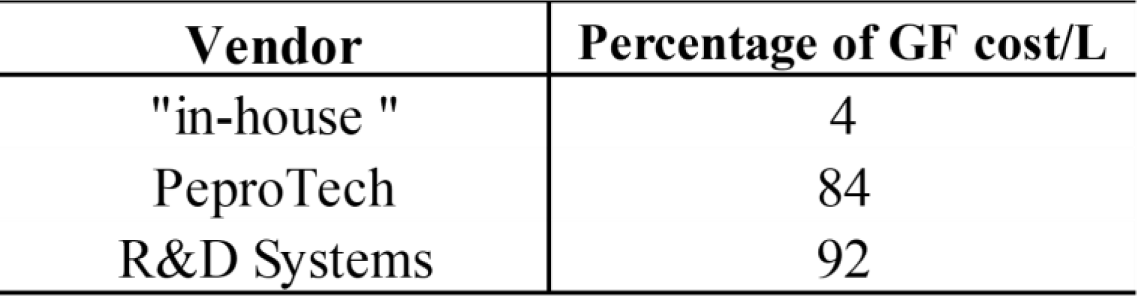
Percentage cost contribution of growth factors (FGF-2 and TGFβ-1) per litre of culture media pricing. DMEM basal media cost per litre is CAD$ 31.06. Growth factors contribute only 4% of the total cost of culture media using our “in-house” purified growth factors.

Beyond the cost of producing the individual GFs, our discovery of specific non-mammalian orthologs that provide comparable or improved bioactivity at lower concentrations also allows for a further reduction in the overall cost of growth medium. Specifically, the ability to achieve comparable cell proliferation with reduced amounts (ng/mL) of FGF-2 (10-fold reduction as in FGF-2 *Atlantic salmon* ortholog) can result in even further reductions in media costs.

## Discussion

The presence of mitogenic GFs is a primary requirement for cell culture medium essential for various applications both in academia and bioindustry including the emerging fields of tissue engineering and cellular agriculture. To address the need for cost effective serum-free eukaryotic cell culture medium, we pursued the development of protocols for producing scalable, low-cost recombinantly expressed GFs commonly used in medium compositions. This new data represents a significant advancement for facilitating cost reductions in basic tissue engineering research applications as well as the large-scale applications anticipated for future CCM production.

Environmental and ethical concerns surrounding traditional animal agriculture have brought increased focus on the development of alternative sources for dietary protein. CCM represents a promising approach for augmenting existing food production in a land, water, and carbon efficient manner while eliminating the need to harvest from whole animals. Despite its promise, CCM production is currently challenged by the high cost associated with *in vitro* cell culture, in particular the costs associated with growth medium. While SFM is effective in supporting *in vitro* cell culture, its cost is often prohibitive for large scale applications, and this is driven primarily by the cost of mitogenic GF components. Addressing this challenge, we have developed an array of expression protocols that facilitate high levels of soluble bioactive GF protein production in *E. coli*, a tractable host that can be cultured with minimal specialized equipment and in BSL1 laboratory facilities.

Our methodology outlined is standardized and based on classic IMAC affinity chromatography techniques, allowing for the same procedure to be used in preparation of bioactive FGF-1 and FGF-2, IGF-1 and IGF-2, PDGF-BB and TGF-β1. The modest demands for laboratory equipment and technical expertise allow for this methodology to be suitable for both academic and industrial settings with minimal equipment and experience in recombinant protein production technology that should facilitate significant reductions in the costs associated with procuring bioactive GFs (Figure 5). Furthermore, we have demonstrated that many of the GFs retain bioactivity as fusion proteins, allowing for purification to proceed via a single chromatography step and without the need for fusion partner cleavage and separation. Despite this observation, we have also outlined and validated strategies for efficient removal of fusion partner tags should it be deemed necessary by end users of these expression systems (Figure S3). To our knowledge, the expression systems we have developed represent the first reported instance of successful expression of soluble and bioactive PDGF-BB and TGF-β1 in *E. coli* expression system.

**Figure 5.**
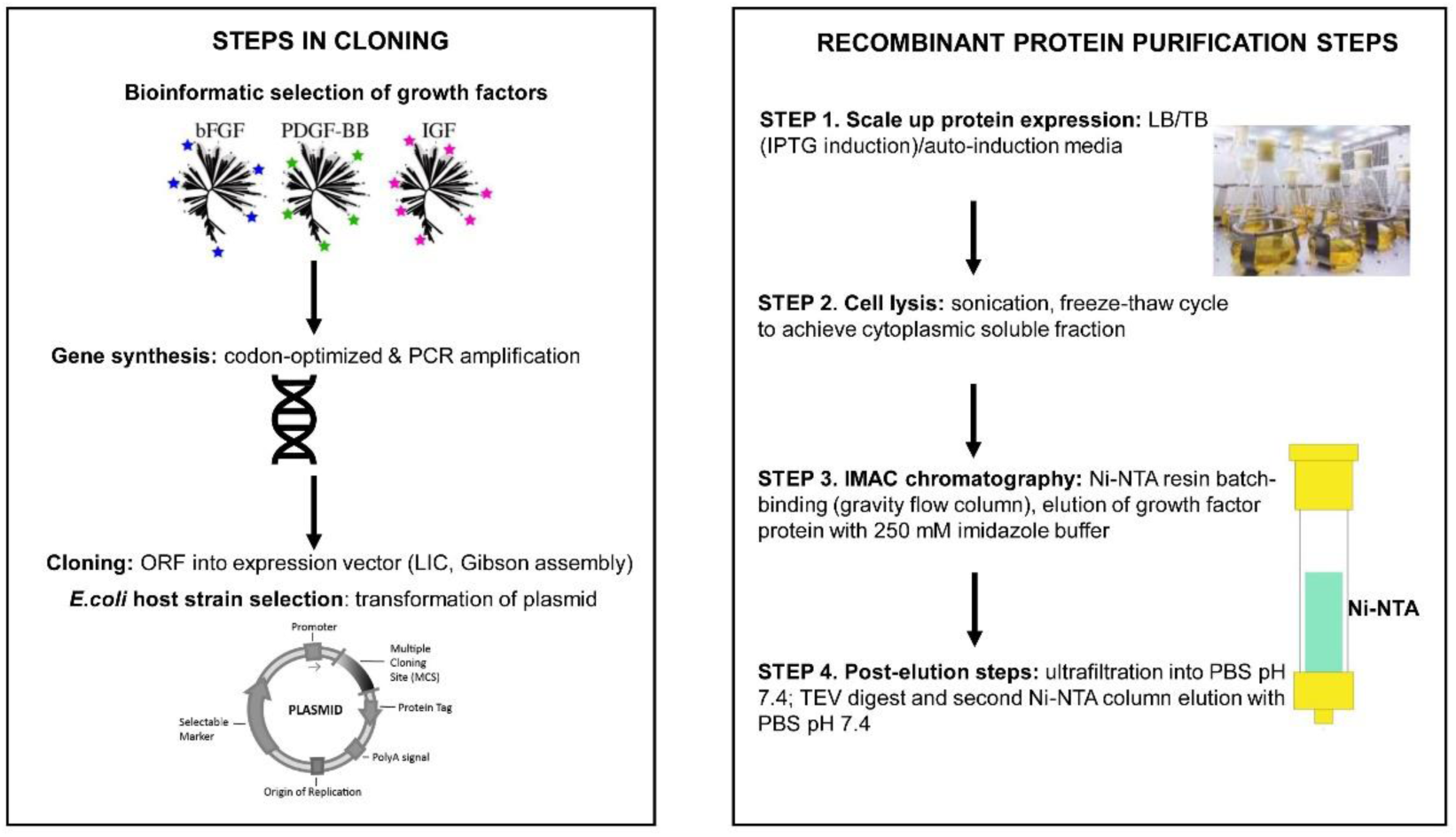
Protein expression and purification strategy followed for the GF proteins. The flowchart specifically highlights the ease of purification steps by IMAC affinity chromatography, requiring the use of Ni-NTA resin and a gravity column.

The presented approach allowed for expression of multiple orthologs from each of the GF families, including sequences from mammalian, avian, and *Teleost* fish origin in accordance with the CCM industry focus on growing skeletal muscle cells belonging to these groups. Though we did not investigate or rationalize why sequence variations amongst GF orthologs dramatically affects their purification and bioactivity characteristics, the ultimate outcome is projected to be a significant cost reduction as decreased amounts of FGF-2, PDGF-BB or TGFβ−1 are suitable for promoting similar fold cell proliferation as the commercial GFs. Further exploration of existing amino acid sequence diversity within these GF families is warranted and may uncover variants that provide improved performance for meeting the unique needs of CCM production.

In screening GF orthologs for bioactivity, we found several FGF-2 orthologs that induced higher levels of cell proliferation on NIH-3T3 cells compared to commercial mammalian FGF-2. This effect was observed not only at the standard working concentration of FGF-2 (10 ng/mL) but also at reduced concentrations of 0.1-1 ng/mL, raising the possibility that FGF-2 orthologs may provide improved proliferation while using less material. Our functional bioactivity results thus prove a significant cost reduction strategy by using decreased concentrations of FGF-2, PDGF-BB or TGF-β1 to promote similar fold cell proliferation as the commercially available GFs.

Analysis of the cost of production showed that GFs generated using the protocols we have outlined provide substantial cost savings compared to commercially available alternatives, significantly reducing the cost contribution of GFs to approximately 4% of the culture medium. Of note, our pricing is based on yields obtained using a bench-scale shake flask approach. It can reasonably be assumed that optimizing growth conditions and transitioning to large-scale bioreactor fermentation could further improve GF yields and facilitate greater cost reductions.

The methodology outlined for soluble expression of key GF components is designed to allow for academic laboratories as well as small-scale cellular agriculture start-ups to bring GF production “in-house”, thereby allowing for significant cost savings. Most of the expression protocols presented in our study are based on a single IMAC purification and require no *in vitro* protein refolding steps. While we have documented additional steps that can improve GF purity (e.g., TEV cleavage, size exclusion chromatography, anion-exchange chromatography), in many cases these are optional and may not be required for GFs aimed to be used in cell culture medium. Since the initial purification steps for all these GFs are identical, a standardized workflow that requires minimal specialized equipment or technical expertise can be adopted to facilitate production of the major cost drivers of SFM (Figure 5).

The expression systems we have developed proved to be robust for a variety of mitogenic GFs and represents a key advancement to meeting the increasing demands for these critical components in cell culture media. The set of expression vectors generated in this study also provide a toolset for bacterial expression of other GF families not tested here, and in general, for disulfide-rich proteins more broadly that may have been recalcitrant to bacterial expression system.

## Limitations of the study

The functional bioactivity of the recombinant growth factors was assayed using fibroblast cell lines. Primary skeletal muscle cells will be a more appropriate model system to test the suitability of their use in cellular agriculture. It was beyond the scope of the work presented to investigate the molecular mechanisms behind the variation in functional activity observed between different growth factor orthologs.

## Resource Availability

### Lead Contact

Further information and requests for resources should be directed to the corresponding authors, Alexei Savchenko and Peter Stogios

### Materials Availability

Plasmids used in this study are available upon request.

## Acknowledgements

We thank Dr. Alison McGuigan and members of the McGuigan lab for allowing us to use their cell culture facility and for helpful input on methodology. We also thank Dr. Tom Ben-Arye, Senior Scientist, The Good Food Institute (GFI) for scientific input and advice. This work has been funded by a research grant from GFI awarded to Peter J Stogios, a New Harvest Postdoctoral Fellowship Award awarded to Cameron Semper, and a grant from the Animal Advocacy Research Fund awarded to Cameron Semper.

## Author Contributions

Conceptualization, M.V., C.S, P.J.S. and A.S.; Methodology, M.V. and C.S.; Investigation, M.V., C.S., R.D. and N.M.; Writing – Original Draft, M.V. and C.S.; Writing – Review and Editing, M.V., C.S., P.J.S. and A.S.; Funding Acquisition – C.S., P.J.S. and A.S.

## Materials and Methods

### Bacterial strains, vector construction

*E.coli* DH5α were used for cloning and vector construction. For protein expression, BL21 (DE3) Gold, Shuffle T7 express (NEB), Origami B (DE3), Rosetta Gami B (DE3) (EMD-Millipore) competent cells were used. Commercially used vectors include pMCSG53-His6x-TEV-LIC (MCSG) with LIC cloning sites, pET-His6x-TrxA-TEV -LIC (Addgene 29712), pET-His6x-SUMO-TEV-LIC (Addgene 29711). The vectors pMCSG53-His6x-DsbC-TEV and pMCSG53-His6x-DsbA-TEV were designed “in-house” using the pMCSG53-His6x-TEV-LIC backbone and introducing the DsbC/DsbA gene in frame downstream of His6x tag.

All the insert gene sequences were codon optimized for *E.coli*expression system and synthesized from Twist BioSciences or Codex DNA. The insert genes were cloned into the vector pMCSG53 or pET-TrxA or pET-SUMO or pMCSG53-DsbC, pMCSG53-DsbA, coding for a fusion protein with an N-terminal 6x histidine tag cleavable by TEV protease. DNA sequencing in the cloned plasmid was verified at the TCAG DNA sequencing facility, SickKids, Toronto. *E.coli*competent cells were transformed with the plasmids. 20 mL of overnight culture (16h approx.) in Luria Bertani (LB) broth was diluted into 1 L of LB or terrific broth (TB) containing selected antibiotics (100 ug/mL ampicillin) and grown at 37 °C using the shaker flask approach. Expression was induced with 0.6 mM IPTG at 17 °C when optical density, OD600 reached 0.8-0.9 absorbance units and allowed to grow overnight for 16-18h. The large overnight cell cultures were then collected by centrifugation at 7000g.

Cells were resuspended in binding buffer [pH 7.5, 100 mM HEPES, 500 mM NaCl, 5 mM imidazole, and 5% glycerol (v/v)] and lysed. The cell debris was removed by centrifugation at 20,000g. Ni-NTA affnity chromatography was used for protein purification. The soluble cytoplasmic fraction containing the GF protein was purified by batch-binding to a Ni-NTA resin, washed with wash buffer [pH 7.5, 100 mM HEPES, 500 mM NaCl, 30 mM imidazole, and 5% glycerol (v/ v)], and the protein was eluted with elution buffer [pH 7.5, 100 mM HEPES, 500 mM NaCl, 250 mM imidazole, and 5% glycerol (v/v)]. The His6-tagged protein was then subjected to overnight (16-24h approx.) TEV protease digest using 50 μg of “in house” TEV protease per mg of His6x-tagged protein and simultaneously dialyzed overnight against phosphate-buffered saline (PBS), pH 7.4 buffer containing no imidazole. The 6xHis-tag and TEV was removed by binding the protein onto a Ni-NTA column again, whereby the flow through contained the His-tag cleaved protein of interest. For plasmid constructs where the TEV protease digest step was skipped, the final proteins were dialyzed with a minimum of 2 x 2L dilution cycles in (PBS) pH 7.4. For PDGF-BB, the His6x-Trx tag was separated from PDGF-BB using size-exclusion chromatography (Superdex 75 16/60). Fractions containing the PDGF-BB were pooled and concentrated. For TGFB-1 and IGF-1/IGF-2, the His6x-DsbC tag was separated from the GF using anion exchange chromatography (MonoQ 5/5). The purity of all the recombinantly purified proteins were analyzed by SDS-polyacrylamide gel electrophoresis. Non-reducing gel electrophoresis was used to confirm the presence of dimeric state of the proteins. The purified proteins were also subjected to size exclusion chromatography (Superdex 200 16/60) analysis for determination of their oligomeric state. The proteins were concentrated using a Vivaspin concentrator (GE Healthcare) and passed through a 0.2 µm Ultrafree-MC centrifugal filter (EMD Millipore) before storing in aliquots at -80 °C.

### Cell culture

NIH 3T3 mouse fibroblast cell lines (ATCC CRL-1658) were cultured in 100 x 20 mm and 150 x 25 mm culture dishes (Falcon Corning cat. no. 353003 and 353025) using Dulbecco’s modified Eagle’s medium containing 4.5 g/L D-glucose, L-glutamine, 110 mg/L sodium pyruvate (DMEM, GIBCO 11995-065) with 10% heat inactivated newborn calf serum, NBCS (GIBCO 26010-074). Primary pancreatic stellate cells (PSC) were cultured until passage 8 in DMEM with 10% heat inactivated fetal bovine serum and penicillin-streptomycin. Cultures were allowed to reach 80% confluency before passaging them into new media. Briefly, cells were washed twice with phosphate-buffered saline (PBS) without calcium and magnesium, Wisent 311-010-CL, cells detached using Trypsin-0.25% EDTA (Wisent) and split at a seeding density of 5,000 cells/cm^2^. Commercial GFs used in this study included the following: human FGF-1 (PeproTech, cat. no. 100-17A), human FGF-2 (PeproTech, cat. no. 100-18B), human PDGF-BB (PeproTech, cat. no. 100-14B), human IGF-1 (PeproTech, cat. no. 100-11), human IGF-2 (PeproTech, cat. no. 100-12), human TGF-β1(PeproTech, cat. no. 100-21C).

### Bioactivity MTT assay

NIH-3T3 cells were cultured in 96-well plates at a seeding density of 6000 cells/well and allowed to attach for 24h. Cells were subsequently serum starved for 24h in 0.11% serum medium. GFs were added to individual wells at specific serial dilution concentrations and incubated at 37 °C with 5% CO2. After 48h, the metabolic activity of viable cells was tested by a colorimetric dimethyl-thiazole-diphenyltetrazolium bromide (MTT) assay (cat. no. M6494, Invitrogen). Briefly, the steps involved aspirating the culture media, washing the wells with PBS to remove any dead cells, and then adding 100 µL of DMEM media containing 10 μL MTT (5 mg/mL in sterile PBS) into each well. The 96 well plates were incubated at 37 °C for 4h. The media was then aspirated, wells washed with 100 μL PBS and the insoluble MTT crystals were solubilized with 50 µL of DMSO by incubating in an orbital plate shaker at 700 rpm, 37 °C for 20-30 min. Absorbance was recorded immediately on a Tecan plate reader at a wavelength of 540 nm. Three independent experiments were conducted for each set of GFs with different passage number cells. Triplicate readings from each set were averaged and data was presented as mean ± SD (SD = standard deviation) using Graphpad Prism v5.0.

### Western blotting

Cells were cultured in 6-well plates at a seeding density of 100,000 cells/well. After 24h, culture medium in the wells was changed into 0.11% serum. Cells were serum starved for 24h before adding the respective GFs. Cell lysis was done after 45 min for TGFβ1, 24h for FGF2, PDGFBB, IGF-1. To extract the total protein, cells were lysed by washing the cells twice in ice-cold PBS, then lysed with 1X RIPA buffer (Cell Signalling Tech. cat. no. 9806), incubated at 4 °C for 1h and centrifuged at 14,000 rpm for 20 min at 4 °C. The supernatant was mixed with 1X SDS-sample buffer containing β-mercaptoethanol, boiled at 95 °C for 10 min and samples loaded onto a 15% SDS-PAGE protein gel. Proteins were transferred to a nitrocellulose membrane using Trans-Blot apparatus (Bio-Rad). After blocking the membrane with 5% w/v BSA and 0.1% v/v Tween-20 in TBS (Tris buffered-saline) for 1h, the membrane was washed 5x with TBST (Tris buffered-saline containing 0.1% v/v Tween-20). The membrane was incubated with the following primary antibodies overnight at 4 °C on a rotating shaker: phospho-smad2 rabbit mAb (Cell Signalling Tech. cat. no. 3108; 1:1000), smad2 rabbit mAb (Cell Signalling Tech. cat. no. 5339; 1:1000), phospho-p44/42 MAPK ERK1/2 rabbit mAb (Cell Signalling Tech., cat. no. 9101; 1:1000), p44/42 MAPK ERK1/2 rabbit mAb (Cell Signalling Tech., cat. no. 4695; 1:1000). The membrane was then incubated with Anti-rabbit IgG, HRP-conjugated secondary antibody (Cell Signalling Tech., cat. no. 7074; 1:2000) with gentle agitation for 1h at room temperature. Immunoactivity was detected using Bio-Rad Clarity Western ECL peroxide-enhancer chemiluminescence kit (cat. no. 170-5060). Chemiluminescent bands were visualized with a Chemidoc XRS+ system (BioRad).

## REFERENCES

Ai, G., Shao, X., Meng, M., Song, L., Qiu, J., Wu, Y., . . . Tong, X. (2017). Epidermal growth factor promotes proliferation and maintains multipotency of continuous cultured adipose stem cells via activating STAT signal pathway in vitro. Medicine (Baltimore), 96(30), e7607. doi:10.1097/MD.0000000000007607

Albrecht, D. E., & Tidball, J. G. (1997). Platelet-derived growth factor-stimulated secretion of basement membrane proteins by skeletal muscle occurs by tyrosine kinase-dependent and -independent pathways. J Biol Chem, 272(4), 2236–2244. doi:10.1074/jbc.272.4.2236

Alexander, D. M., Hesson, T., Mannarino, A., Cable, M., & Dalie, B. L. (1992). Isolation and purification of a biologically active human platelet-derived growth factor BB expressed in Escherichia coli. Protein Expr Purif, 3(3), 204–211. doi:10.1016/1046-5928(92)90016-p

Benington, L., Rajan, G., Locher, C., & Lim, L. Y. (2020). Fibroblast Growth Factor 2-A Review of Stabilisation Approaches for Clinical Applications. Pharmaceutics, 12(6). doi:10.3390/pharmaceutics12060508

Bessette, P. H., Aslund, F., Beckwith, J., & Georgiou, G. (1999). Efficient folding of proteins with multiple disulfide bonds in the Escherichia coli cytoplasm. Proc Natl Acad Sci U S A, 96(24), 13703–13708. doi:10.1073/pnas.96.24.13703

Biancucci, M., Dolores, J. S., Wong, J., Grimshaw, S., Anderson, W. F., Satchell, K. J., & Kwon, K. (2017). New ligation independent cloning vectors for expression of recombinant proteins with a self-cleaving CPD/6xHis-tag. BMC Biotechnol, 17(1), 1. doi:10.1186/s12896-016-0323-4

Boudreault, P., Tremblay, J. P., Pepin, M. F., & Garnier, A. (2001). Scale-up of a myoblast culture process. J Biotechnol, 91(1), 63–74. doi:10.1016/s0168-1656(01)00291-7

Breitkopf, K., Roeyen, C., Sawitza, I., Wickert, L., Floege, J., & Gressner, A. M. (2005). Expression patterns of PDGF-A, -B, -C and -D and the PDGF-receptors alpha and beta in activated rat hepatic stellate cells (HSC). Cytokine, 31(5), 349–357. doi:10.1016/j.cyto.2005.06.005

Brewer, J. R., Mazot, P., & Soriano, P. (2016). Genetic insights into the mechanisms of Fgf signaling. Genes Dev, 30(7), 751–771. doi:10.1101/gad.277137.115

Brown, J., Delaine, C., Zaccheo, O. J., Siebold, C., Gilbert, R. J., van Boxel, G., . . . Jones, E. Y. (2008). Structure and functional analysis of the IGF-II/IGF2R interaction. EMBO J, 27(1), 265–276. doi:10.1038/sj.emboj.7601938

Chen, G., Gulbranson, D. R., Hou, Z., Bolin, J. M., Ruotti, V., Probasco, M. D., . . . Thomson, J. A. (2011). Chemically defined conditions for human iPSC derivation and culture. Nat Methods, 8(5), 424–429. doi:10.1038/nmeth.1593

Cheng, Y., & Patel, D. J. (2004). An efficient system for small protein expression and refolding. Biochem Biophys Res Commun, 317(2), 401–405. doi:10.1016/j.bbrc.2004.03.068

Derynck, R., Jarrett, J. A., Chen, E. Y., Eaton, D. H., Bell, J. R., Assoian, R. K., . . . Goeddel, D. V. (1985). Human transforming growth factor-beta complementary DNA sequence and expression in normal and transformed cells. Nature, 316(6030), 701–705. doi:10.1038/316701a0

Emamipour, N., Vossoughi, M., Mahboudi, F., Golkar, M., & Fard-Esfahani, P. (2019). Soluble expression of IGF1 fused to DsbA in SHuffle T7 strain: optimization of expression and purification by Box-Behnken design. Appl Microbiol Biotechnol, 103(8), 3393–3406. doi:10.1007/s00253-019-09719-w

Eschenfeldt, W. H., Makowska-Grzyska, M., Stols, L., Donnelly, M. I., Jedrzejczak, R., & Joachimiak, A. (2013). New LIC vectors for production of proteins from genes containing rare codons. J Struct Funct Genomics, 14(4), 135–144. doi:10.1007/s10969-013-9163-9

Fredriksson, L., Li, H., & Eriksson, U. (2004). The PDGF family: four gene products form five dimeric isoforms. Cytokine Growth Factor Rev, 15(4), 197–204. doi:10.1016/j.cytogfr.2004.03.007

Gasparian, M. E., Elistratov, P. A., Drize, N. I., Nifontova, I. N., Dolgikh, D. A., & Kirpichnikov, M. P. (2009). Overexpression in Escherichia coli and purification of human fibroblast growth factor (FGF-2). Biochemistry (Mosc), 74(2), 221–225. doi:10.1134/s000629790902014x

Geistlinger, T., Jhala, R., Krueger, K., Ramesh, B. (2017). United States Patent No.: W. I. P. Organization.

Godfray, H. C. J., Aveyard, P., Garnett, T., Hall, J. W., Key, T. J., Lorimer, J., . . . Jebb, S. A. (2018). Meat consumption, health, and the environment. Science, 361(6399). doi:10.1126/science.aam5324

Hakuno, F., & Takahashi, S. I. (2018). IGF1 receptor signaling pathways. J Mol Endocrinol, 61(1), T69–T86. doi:10.1530/JME-17-0311

Hinck, A. P., Archer, S. J., Qian, S. W., Roberts, A. B., Sporn, M. B., Weatherbee, J. A., . . . Torchia, D. A. (1996). Transforming growth factor beta 1: three-dimensional structure in solution and comparison with the X-ray structure of transforming growth factor beta 2. Biochemistry, 35(26), 8517–8534. doi:10.1021/bi9604946

Kim, Y. S., Lee, H. J., Han, M. H., Yoon, N. K., Kim, Y. C., & Ahn, J. (2021). Effective production of human growth factors in Escherichia coli by fusing with small protein 6HFh8. Microb Cell Fact, 20(1), 9. doi:10.1186/s12934-020-01502-1

Koledova, Z., Sumbal, J., Rabata, A., de La Bourdonnaye, G., Chaloupkova, R., Hrdlickova, B., . . . Stepankova, V. (2019). Fibroblast Growth Factor 2 Protein Stability Provides Decreased Dependence on Heparin for Induction of FGFR Signaling and Alters ERK Signaling Dynamics. Front Cell Dev Biol, 7, 331. doi:10.3389/fcell.2019.00331

Kuo, H. H., Gao, X., DeKeyser, J. M., Fetterman, K. A., Pinheiro, E. A., Weddle, C. J., . . . Burridge, P. W. (2020). Negligible-Cost and Weekend-Free Chemically Defined Human iPSC Culture. Stem Cell Reports, 14(2), 256–270. doi:10.1016/j.stemcr.2019.12.007

LaVallie, E. R., DiBlasio, E. A., Kovacic, S., Grant, K. L., Schendel, P. F., & McCoy, J. M. (1993). A thioredoxin gene fusion expression system that circumvents inclusion body formation in the E. coli cytoplasm. Biotechnology (N Y), 11(2), 187–193. doi:10.1038/nbt0293-187

Lobstein, J., Emrich, C. A., Jeans, C., Faulkner, M., Riggs, P., & Berkmen, M. (2012). SHuffle, a novel Escherichia coli protein expression strain capable of correctly folding disulfide bonded proteins in its cytoplasm. Microb Cell Fact, 11, 56. doi:10.1186/1475-2859-11-56

Mattick, C. S., Landis, A. E., Allenby, B. R., & Genovese, N. J. (2015). Anticipatory Life Cycle Analysis of In Vitro Biomass Cultivation for Cultured Meat Production in the United States. Environ Sci Technol, 49(19), 11941–11949. doi:10.1021/acs.est.5b01614

McCubrey, J. A., Steelman, L. S., Mayo, M. W., Algate, P. A., Dellow, R. A., & Kaleko, M. (1991). Growth-promoting effects of insulin-like growth factor-1 (IGF-1) on hematopoietic cells: overexpression of introduced IGF-1 receptor abrogates interleukin-3 dependency of murine factor-dependent cells by a ligand-dependent mechanism. Blood, 78(4), 921–929.

Melzener, L., Verzijden, K. E., Buijs, A. J., Post, M. J., & Flack, J. E. (2021). Cultured beef: from small biopsy to substantial quantity. J Sci Food Agric, 101(1), 7–14. doi:10.1002/jsfa.10663

Mossahebi-Mohammadi, M., Quan, M., Zhang, J. S., & Li, X. (2020). FGF Signaling Pathway: A Key Regulator of Stem Cell Pluripotency. Front Cell Dev Biol, 8, 79. doi:10.3389/fcell.2020.00079

Mullen, A. C., & Wrana, J. L. (2017). TGF-beta Family Signaling in Embryonic and Somatic Stem-Cell Renewal and Differentiation. Cold Spring Harb Perspect Biol, 9(7). doi:10.1101/cshperspect.a022186

Nozach, H., Fruchart-Gaillard, C., Fenaille, F., Beau, F., Ramos, O. H., Douzi, B., . . . Dive, V. (2013). High throughput screening identifies disulfide isomerase DsbC as a very efficient partner for recombinant expression of small disulfide-rich proteins in E. coli. Microb Cell Fact, 12, 37. doi:10.1186/1475-2859-12-37

Peruzzi, F., Prisco, M., Dews, M., Salomoni, P., Grassilli, E., Romano, G., . . . Baserga, R. (1999). Multiple signaling pathways of the insulin-like growth factor 1 receptor in protection from apoptosis. Mol Cell Biol, 19(10), 7203–7215. doi:10.1128/MCB.19.10.7203

Poniatowski, L. A., Wojdasiewicz, P., Gasik, R., & Szukiewicz, D. (2015). Transforming growth factor Beta family: insight into the role of growth factors in regulation of fracture healing biology and potential clinical applications. Mediators Inflamm, 2015, 137823. doi:10.1155/2015/137823

Post, M. J., Levenberg, S., Kaplan, D. L., Genovese, N., Fu, J., Bryant, C. J., . . . Moutsatsou, P. (2020). Scientific, sustainability and regulatory challenges of cultured meat. Nature Food, 1(7), 403–415. doi:10.1038/s43016-020-0112-z

Rosano, G. L., & Ceccarelli, E. A. (2014). Recombinant protein expression in Escherichia coli: advances and challenges. Front Microbiol, 5, 172. doi:10.3389/fmicb.2014.00172

Schafer, F., Seip, N., Maertens, B., Block, H., & Kubicek, J. (2015). Purification of GST-Tagged Proteins. Methods Enzymol, 559, 127–139. doi:10.1016/bs.mie.2014.11.005

Scharenberg, M. A., Pippenger, B. E., Sack, R., Zingg, D., Ferralli, J., Schenk, S., . . . Chiquet-Ehrismann, R. (2014). TGF-beta-induced differentiation into myofibroblasts involves specific regulation of two MKL1 isoforms. J Cell Sci, 127(Pt 5), 1079–1091. doi:10.1242/jcs.142075

Schmierer, B., & Hill, C. S. (2007). TGFbeta-SMAD signal transduction: molecular specificity and functional flexibility. Nat Rev Mol Cell Biol, 8(12), 970–982. doi:10.1038/nrm2297

Spangenburg, E. E., & Booth, F. W. (2002). Multiple signaling pathways mediate LIF-induced skeletal muscle satellite cell proliferation. Am J Physiol Cell Physiol, 283(1), C204–211. doi:10.1152/ajpcell.00574.2001

Specht, L. (2020). An analysis of culture medium costs and production volumes for cultivated meat Retrieved from https://gfi.org/wp-content/uploads/2021/01/clean-meat-production-volume-and-medium-cost.pdf:

Stout, A. J., Mirliani, A. B., White, E. C., Yuen, J. S. K., & Kaplan, D. L. (2021). Simple and effective serum-free medium for sustained expansion of bovine satellite cells for cell cultured meat. bioRxiv, 2021.2005.2028.446057. doi:10.1101/2021.05.28.446057

Tripathi, N. K., & Shrivastava, A. (2019). Recent Developments in Bioprocessing of Recombinant Proteins: Expression Hosts and Process Development. Front Bioeng Biotechnol, 7, 420. doi:10.3389/fbioe.2019.00420

Vajdos, F. F., Ultsch, M., Schaffer, M. L., Deshayes, K. D., Liu, J., Skelton, N. J., & de Vos, A. M. (2001). Crystal structure of human insulin-like growth factor-1: detergent binding inhibits binding protein interactions. Biochemistry, 40(37), 11022–11029. doi:10.1021/bi0109111

van der Valk, J., Bieback, K., Buta, C., Cochrane, B., Dirks, W. G., Fu, J., . . . Gstraunthaler, G. (2018). Fetal Bovine Serum (FBS): Past - Present - Future. ALTEX, 35(1), 99–118. doi:10.14573/altex.1705101

Yang, Z., & Xiong, H.-R. (2012). Culture Conditions and Types of Growth Media for Mammalian Cells. In L. Ceccherini-Nelli & B. Matteoli (Eds.), Biomedical Tissue Culture. DOI: 10.5772/52301: IntechOpen.

Yao, T., & Asayama, Y. (2017). Animal-cell culture media: History, characteristics, and current issues. Reprod Med Biol, 16(2), 99–117. doi:10.1002/rmb2.12024

Yu, M., Wang, H., Xu, Y., Yu, D., Li, D., Liu, X., & Du, W. (2015). Insulin-like growth factor-1 (IGF-1) promotes myoblast proliferation and skeletal muscle growth of embryonic chickens via the PI3K/Akt signalling pathway. Cell Biol Int, 39(8), 910–922. doi:10.1002/cbin.10466

Zakrzewska, M., Krowarsch, D., Wiedlocha, A., & Otlewski, J. (2004). Design of fully active FGF-1 variants with increased stability. Protein Eng Des Sel, 17(8), 603–611. doi:10.1093/protein/gzh076

Zheng, W., et al (2016). Expression and purification of human epidermal growth factor (hEGF) fused with GB1. Biotechnology & Biotechnological Equipment, 30(4), 813–818.

Zhu, H., Duchesne, L., Rudland, P. S., & Fernig, D. G. (2010). The heparan sulfate co-receptor and the concentration of fibroblast growth factor-2 independently elicit different signalling patterns from the fibroblast growth factor receptor. Cell Commun Signal, 8, 14. doi:10.1186/1478-811X-8-14

Zou, Z., & Sun, P. D. (2004). Overexpression of human transforming growth factor-beta1 using a recombinant CHO cell expression system. Protein Expr Purif, 37(2), 265–272. doi:10.1016/j.pep.2003.06.001

