## Supplemental Info for "Recombinant production of growth factors for application in cell culture"

### SUPPLEMENTARY METHODS

#### Additional protein purification steps for growth factors:

Some PDGF-BB orthologs required an additional step after the second Ni-NTA step. The cut PDGF-BB and the Trx tag co-eluted and needed an additional size-exclusion chromatography (SEC) step. The eluent from the Ni-NTA was applied to Superdex 75 16/60 column (AKTA systems) using PBS pH 7.4 at a flow rate of 1 mL/min and 0.5 mL fractions were collected over 1.5x column volume elution. Fractions were analyzed on SDS-PAGE and PDGF-BB containing fractions were pooled and concentrated.

IGF-1/IGF-2 and TGFB-1 orthologs resulted in co-elution of cut-IGF/TGFb-1 after the TEV digest and second Ni-NTA column elution. The eluent was applied to an anion exchange chromatography (IEX), MonoQ HR 5/5 (AKTA systems) using 50 mM CAPS pH 10.65, 10 mM NaCl (Buffer A) and 50 mM CAPS pH 10.65, 1M NaCl. A stepwise gradient of 30%B resulted in separating the DsbC tag to a significant extent.

### SUPPLEMENTARY FIGURES

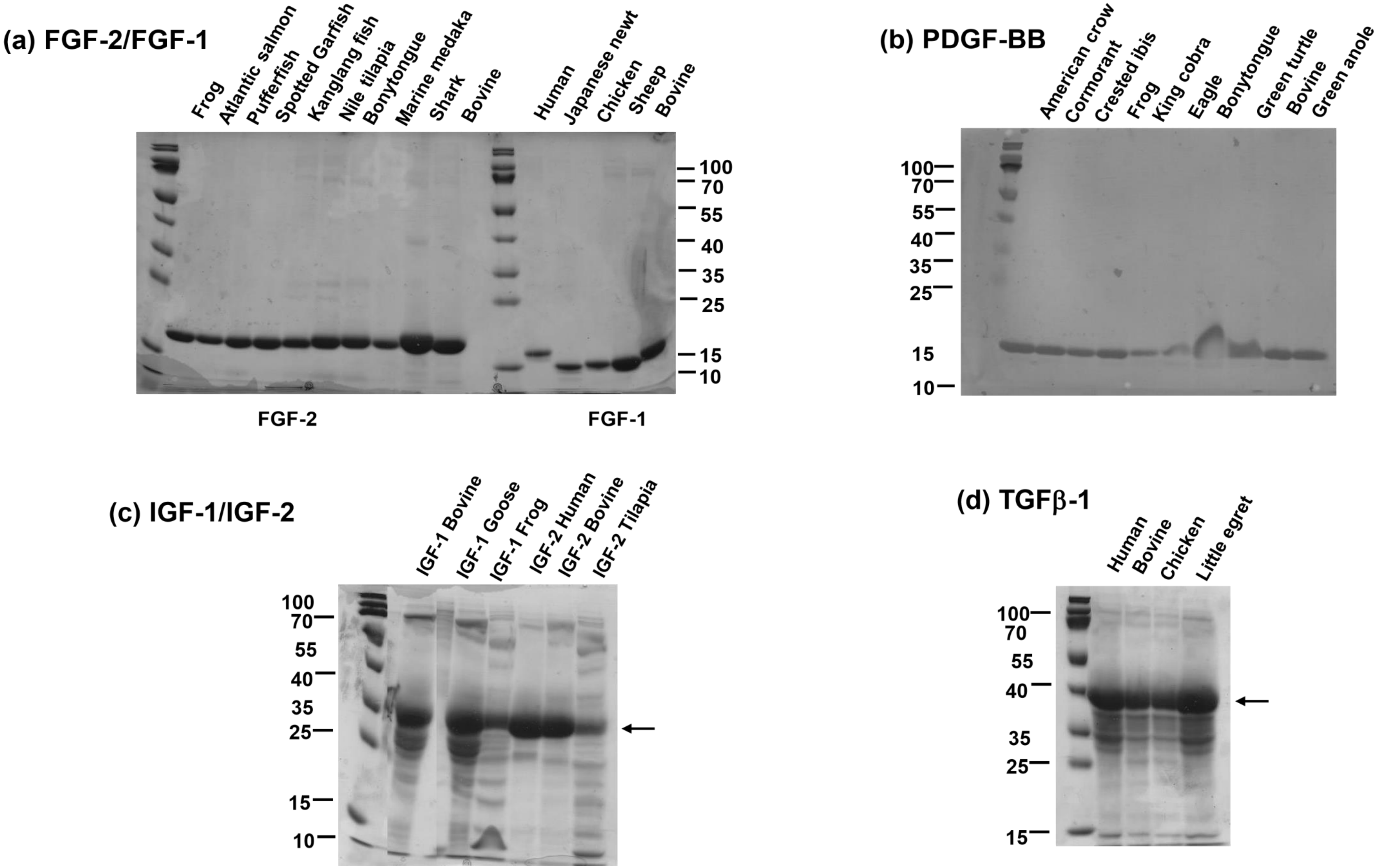

**Supplementary figure 1. Summary of a subset of GF targets recombinantly purified in this study.** Analyzed on a 15% reducing SDS-PAGE gel electrophoresis **(a)** FGF-2 and FGF-1 orthologs, 15 kDa **(b)** PDGF-BB orthologs, 15 kDa **(c)** IGF-1, IGF-2 orthologs, 35 kDa and **(d)** TGFβ-1 orthologs, 40 kDa

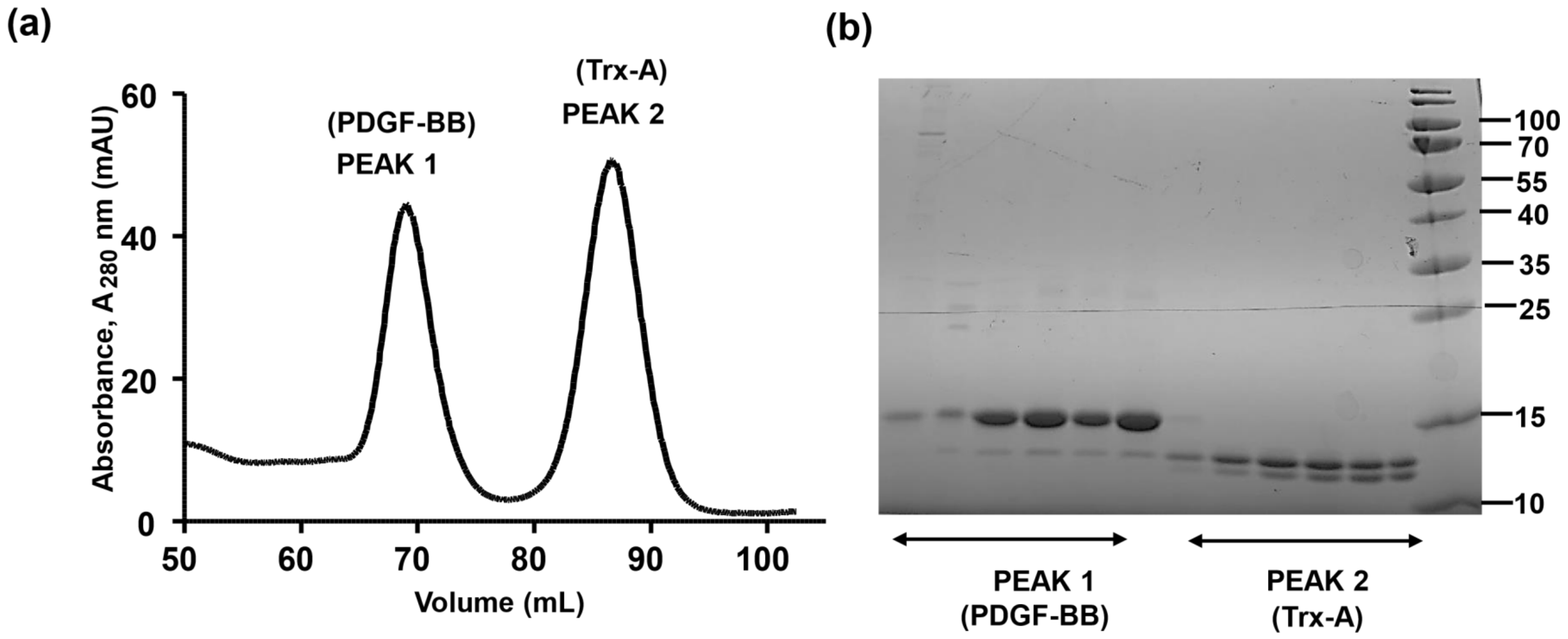

**Supplementary Figure 2**. **(a)**Size-exclusion chromatography Superdex 75 16/60 for separation of PDGF-BB from the TrxA tag after TEV digest. **(b)** Fractions corresponding to **PEAK 1** and **PEAK 2** were analyzed on SDS-PAGE. The fractions corresponding to PDGFBB (**PEAK 1**) was pooled and concentrated.

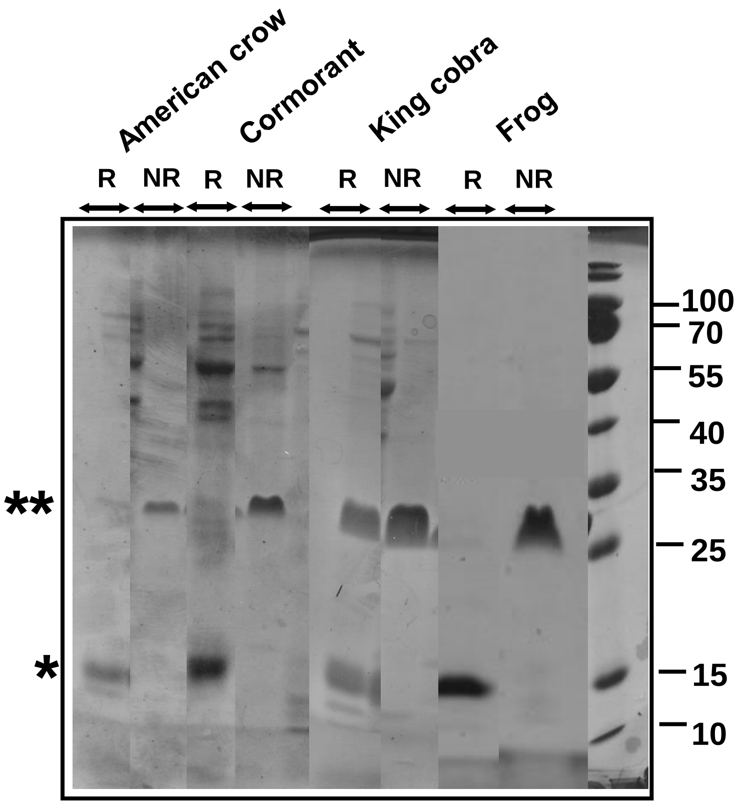

**Supplementary Figure 3.** Recombinantly purified PDGF-BB orthologs analyzed under reducing **(R)** and non-reducing **(NR)** conditions on a 15% gel electrophoresis. Under non-reducing conditions, all PDGF-BB orthologs dimerize marked with double asterik (**) at 25 kDa approx.

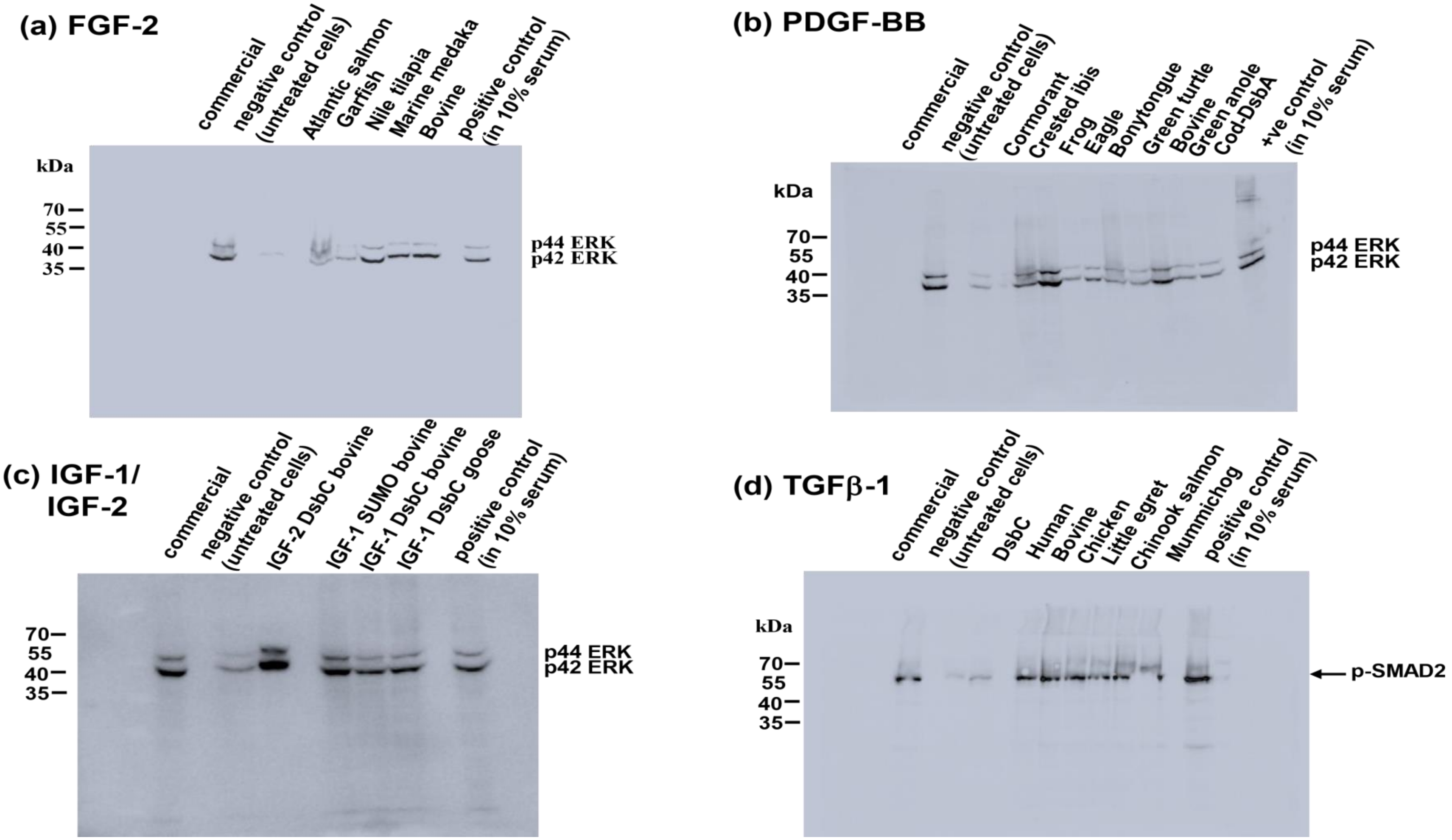

**Supplementary Figure 4.** Western blot nitrocellulose membrane images of **(a)** FGF-2 (p-ERK1/2) **(b)** PDGF-BB (p-ERK1/2) **(c)** IGF-1/IGF-2 (p-ERK1/2) **(d)** TGFβ-1 (p-SMAD2)

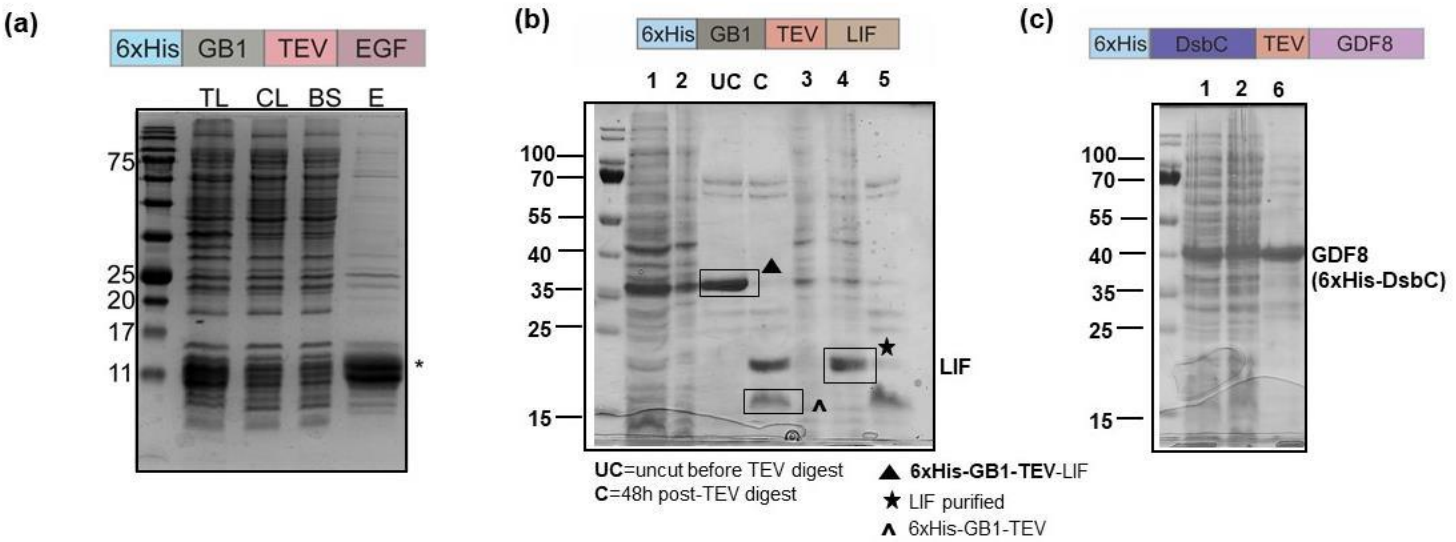

**Supplementary Figure 5.** **(a)** Scale up expression of EGF human orthologue expressed in BL21(DE3) Gold cells **(b)** Scale up expressions of LIF human ortholog expressed in BL21(DE3) Gold cells **(c)** GDF8 bovine ortholog expressed in Shuffle T7 express cells. *E.coli* cells grown in 1L terrific broth (TB), induced with 0.8 mM IPTG at $OD_{600}$ of 1.5-2.0 units. *UC=uncut before TEV digest; C=24h post TEV digest.* After the TEV digest, LIF band can be seen at 20 kDa corresponding to the monomer LIF while the His6x-GB1-tag runs at 15 kDa, **1** – soluble fraction; **2** – supernatant post Ni-NTA batch bind; **3** – flow through after second Ni- binding post-TEV digest; **4** – 30 mM imidazole wash eluent; **5** – 250 mM imidazole wash eluent; **6** – GDF8 bovine ortholog purified post-Ni-NTA batch binding

### SUPPLEMENTARY TABLES

**Supplementary Table 1.** Expression constructs/tags used for soluble protein expression of growth factor targets

| Growth factor family | Expression System | *E.coli* host strain |
|---|---|---|
| aFGF/FGF-1 | pMCSG53-His6-TEV | BL21 DE3 Gold |
| bFGF/FGf-2 | pMCSG53-His6-TEV | BL21 DE3 Gold |
| PDGF-BB | pET-Trx-His6-TEV | SHuffle T7 express |
| IGF-1 | pMCSG53-His6-DsbC-TEV | SHuffle T7 express |
| IGF-2 | pMCSG53-His6-DsbC-TEV | SHuffle T7 express |
| TGFb1 | pMCSG53-His6-DsbC-TEV | SHuffle T7 express |
| GDF8 (myostatin) | pMCSG53-His6-DsbC-TEV | SHuffle T7 express |
| LIF | pMCSG53-His6-GB1-TEV | BL21 DE3 Gold |
| EGF | pMCSG53-His6-GB1-TEV | BL21 DE3 Gold |

**Supplementary Table 2.** Protein sequences of growth factors expressed and purified in this study

| Growth factor | GenBank Acc. No. | Amino acid sequence | Organism |
|---|---|---|---|
| FGF-1 | P05230 | FNLPPGNYKKPKLLYCSNGGHFLRILPDGTVDGTRDRSDQ<br>HIQLQLSAESVGEVYIKSTETGQYLAMDTDGLLYGSQTPN<br>EECLFLERLEENHYNTYISKKHAEKNWFVGLKKNGSCKR<br>GPRTHYGQKAILFLPLPVSSD | *Homo sapiens* (Human) |
| FGF-1 | Q6I6M7 | FNLPNGNYQRPKLLYCSNGGHFLRILPDGKVDGTRDRSDP<br>YIQLQFYAESVGEVYIKSLETGQYLAMDSDGRLYASQSPS<br>EECLFLERLEENHYNTYKSKMHADKDWFVGIKKNGKTKL<br>GSRTHFGQKAILFLPLPV | *Cynops pyrrhogaster* (Japanese newt) |
| FGF-1 | P19596 | FGLPLGNYKKPKLLYCSNGGHFLRILPDGKVDGTRDRSDQ<br>HIQLQLSAEDVGEVYIKSTASGQYLAMDTNGLLYGSQLP<br>GEECLFLERLEENHYNTYISKKHADKNWFVGLKKNGNSK<br>LGPRTHYGQKAILFLPLPVSAD | *Gallus gallus* (Chicken) |
| FGF-1 | Q7M303 | FNLPLGNYKKPKLLYCSNGGYFLRILPDGRVDGTKDRSD<br>QHIQLQLYAESIGEVYIKSTETGQFLAMDTNGLLYGSQTP<br>SEECLFLERLEENHYNTYISKKHAEKNWFIGLKKNGSSKL<br>GPRTHFGQKAILFLPLPVSSD | *Ovis aries* (Sheep) |
| FGF-1 | P03968 | FNLPLGNYKKPKLLYCSNGGYFLRILPDGTVDGTKDRSD<br>QHIQLQLCAESIGEVYIKSTETGQFLAMDTDGLLYGSQTP<br>NEECLFLERLEENHYNTYISKKHAEKHWFVGLKKNGRSKL<br>GPRTHFGQKAILFLPLPVSSD | *Bos taurus* (Bovine) |
| FGF-2 | P12226 | PTESEDGGNTPFSPGSFKDPKRLYCKNGGFFLRINSDGRV<br>DGSRDKSDSHIKLQLQAVERGVVSIKGITANRYLAMKEDG<br>RLTSLRCITDECFFFERLEANNYNTYRSRKYSSWYVALKR<br>TGQYKNGSSTGPGQKAILFLPMSAKS | *Xenopus laevis* (African clawed frog) |
| FGF-2 | A0A1S3RRG7 | PATPEDGGSGGFLPGNFKEPKRLYCKNGGYFLRINSNGS<br>VDGIRDKNDPHNKLQLQATSVGEVVIKGVSANRYLAMN<br>ADGRLFGARRTTDECYFMERLESNNYNTYRSRKYPEMY<br>VALKRTGQYKSGSKTGPGQKAILFLPMSARR | *Salmo salar* (Atlantic salmon) |
| FGF-2 | Q4S3V1 | PSTPEDGGSSGFPPGSFKDPKRLYCKNGGFFLRIKSDGVV<br>DGIREKSDPHIKLQLQATSVGEVVIKGVCANRYLAMNRD<br>GRLFGTKRATDECHFLERLESNNYNTYRSRKYPTMFVGL<br>TRTGQYKSGSKTGPGQKAILFLPMSAKC | *Tetraodon nigroviridis* (Spotted green pufferfish) |
| FGF-2 | W5M4K8 | PPAPEDGSSGTFPPGNFKEPKRLYCKNGGYFLRINSDGRI<br>DGIREKNDPHIKLQLQATAVGEVIIKGIYANRYLAMNEDG<br>RLFGSKCHTDECYFIERLESNNYNTYRSKKYPEWYVALK<br>RTGQFKSGSRTGPGQKAILFLPMSAKS | *Lepisosteus oculatus* (Spotted gar) |
| FGF-2 | A0A3N0Y4L3 | PASPDDGASGGFPSGNFKEPKRLYCKNGGYFLRINSDGR<br>VDGIREKSDPHIRLQLQATAVGEVVIKGICANRYLAMNAD<br>GRLFGSRRTTDECYFLERLESNNYNTYRSRKYPDWYVAL<br>KRTGQYKSGSKTSPGQKAILFLPMTAKC | *Anabarilius grahami* (Kanglang fish) |

| | | | |
|---|---|---|---|
| FGF-2 | I3IYU2 | PATPEDGGSSGFPPGNFKDPKRLYCKNGGFFLRIKSDGGV<br>DGIREKNDPHIKLQLQATSVGEVVIKGICANRYLAMNRDG<br>RLFGARRATDECYFLERLESNNYNTYRSRKYPNMYVALK<br>RTGQYKSGSKTGPGQKAILFLPMSAKC | *Oreochromis niloticus* (Nile tilapia) |
| FGF-2 | A0A0P7TW49 | GGAAAAAPGNFKEPKRLYCKNGGYFLRIHPDGRVDGIRD<br>KSDPHIKLQLQATSVGEVVIKGLSANRYLAMNADGRLFG<br>MRRPTDECYFIERLEANNYNTYRSRKYPNWYVALKRTG<br>QYKSGSKTGPGQKAILFLPMSAKC | *Scleropages formosus* (Asian bonytongue) |
| FGF-2 | A0A3B3D1F6 | PEDSGSDSFPPGNFKDPKRLYCKNGGFFLRIRPDGGVDG<br>VREKKDPHIKLRLQATSAGEVVIKGVCSNRYLAMHGDGR<br>LFGVRQATEECYFLERLESNNYNTYRSRRYPNMYVALKR<br>TGQFKPGNKTGPGQKAILFLPMSAKY | *Oryzias melastigma* (Marine medaka) |
| FGF-2 | V9LF85 | PSLPESDPNPFPPGAFKDPKRLYCKNGGYFLRILPDGRVD<br>GTRERSDTRIKLQLQAETVGVISIKGVCANRYLAMNEDGK<br>LYGSKQTTDECFFHERLEPNNYNTYRSKKYDNWYVALK<br>RSGQYKPGPKTGLGQKAILFLPMSAKCS | *Callorhinchus milii* (Ghost shark) |
| FGF-2 | P03969 | PALPEDGGSGAFPPGHFKDPKRLYCKNGGFFLRIHPDGRV<br>DGVREKSDPHIKLQLQAEERGVVSIKGVCANRYLAMKED<br>GRLLASKCVTDECFFFERLESNNYNTYRSRKYSSWYVAL<br>KRTGQYKLGPKTGPGQKAILFLPMSAKS | *Bos taurus* (Bovine) |

| | | | |
|---|---|---|---|
| FGF-2 | A0A6P6LAK1 | PASPDDGASGGFPSGNFKEPKRLYCKNGGFFLRINSDGRV<br>DGIREKSDPHIRLQLQATAVGEVLIKGICANRYLAMNSDG<br>RLIGTRRTTDECYFLERLESNNYNTYRSRKYPDWYVALK<br>RTGQYKAGSKTSPGQKAILFLPMSAK | *Carassius aurantus* (Goldfish) |
| FGF-2 | A0A0R4IHF9 | PAAPDAENSSFPAGSFRDPKRLYCKNGGFFLRINADGRVD<br>GARDKSDPHIRLQLQATAVGEVLIKGICTNRFLAMNADG<br>RLFGTKRTTDECYFLERLESNNYNTYRSRKYPDWYVALK<br>RTGQYKSGSKTSPGQKAILFLPMSAKC | *Danio rerio* (Zebrafish) |
| FGF-2 | A0A060Y7E5 | PATPEDGGSGGFLPGNFKEPKRLYCKNGGYFLRINSNGSV<br>DGIRDKNDPHNKLQLQATSVGEVVIKGVSANRYLAMNA<br>DGRLFGARRTTDECYFMERLESNNYNTYRSRKYPEMYV<br>ALKRTGQYKSGSKTGPGQKAILFLPMSARR | *Oncorhynchus tshawytcha* (Chinook salmon) |
| FGF-2 | A0A437CYC7 | PSPAENSRSDGFPPGNYKDPKRLYCKNGGLFLRIKPDGGV<br>DGIREKKDPHVKLRLQATSAGEVVIKGVCSNRYLAMHGD<br>GRLFGVRQATEECYFLERLESNNYNTYRSKKYPNMYVAL<br>KRTGQYKPGNKTGPGQKAILFLPMSAKY | *Oryzias latipes* (Japanese ricefish) |
| FGF-2 | XP_023646287.1 | PAAPEDGGSGSFLPGTFKELKKLYCKNGGYFLRINADGR<br>VDGTREKTDTHIKLQIQATSIGVVVIKGVSSSRYLAMNDD<br>GRLFGTKRATDECYFFERLESNNYNTYRSREHPNWYVAL<br>KRTGQYKLGSRTGPGQKAILFLPMSA | *Paramormyrops kingsleyae* (Elephantfish) |

| Growth factor | GenBank Acc. No. | Amino acid sequence | Organism |
|---|---|---|---|
| PDGF-BB | A0A091EXH6 | SLDALAAAETAVLAECKTREVVFEISRNMVDSTNANFVV<br>WPPCVEVQRCSGCCNNRNVQCRPTQIRVRHVQVKKIEFV<br>QRKPKFKNVVVPLEDHVQCRCEAVFR | *Corvus brachyrhynchos* (American crow) |

| | | | |
|---|---|---|---|
| PDGF-BB | A0A093QQC3 | SLDALAAAEPAVLAECKTRVVVFEISRNMVDSTNANFVVWPPCVEVQRCSGCCNNRNVQCRPMQIRVRHVQVNKIEFVQRKPKFKKVIVPLEDHVQCRCEAVSS | *Phalacrocorax carbo* (Great cormorant) |
| PDGF-BB | A0A091W1N1 | SLDALAAAEPAVLAECKTRTVVFEISRNMVDSTNANFVVWPPCVEVQRCSGCCNNRNVQCRPTQIRVRPVQVNKIEFVQRKPKFKKVVVPLEDHVQCRCEAVSR | *Nipponia nippon* (Crested ibis) (Ibis nippon) |
| PDGF-BB | B1H1E3 | SLDAEKAVIAECKPRVEVFEISRKIVDPTNANFLVWPPCVEVQRCSGCCNSKNMRCAPTRIHVRHVQVNKIFITPKGKKQVKVVVPLEDHHDCKCEPVPSS | *Xenopus tropicalis* (Western clawed frog) |
| PDGF-BB | V8NSC5 | SVGGAEPIEPAVIAECKTRSEVFEISRSMVDSTNANFIVWPPCVEVQRCTGCCNTRAMQCRPTQVRVRHIQVNKIEVADKKPVFNKAIVALEDHLACRCEPVSP | *Ophiophagus hannah* (King cobra) |
| PDGF-BB | A0A091PJT8 | SLDALAAAEPAVLAECKTRAVVFEISRNMVDSTNANFVVWPPCVEVQRCSGCCNNRNVQCRPTQIRVRHVQVNKIEFVQRKPKFKKVVVPLEDHVQCRCEAVSR | *Haliaeetus albicilla* (White-tailed sea-eagle) |
| PDGF-BB | A0A0P7TR46 | SLEPQPAQQALCKVRTEVLEVTRTMLDRRNANFLLWPPCVEVQRCSGCCNARTVQCMPTVTHMRYLQVMKIQYVNKQPHYEKAVISVVDHIECRCQIEAPAK | *Scleropages formosus* (Asian bonytongue) |
| PDGF-BB | M7AP33 | AEPAVLAECKTRTEVFEISRSMVDSTNANFVVWPPCVEVQRCSGCCNNRNMQCRPTMVHVRHVQVNKIEFIQKKPIFKKAIVPLEDHLECRCEALSS | *Chelonia mydas (*Green sea-turtle) |
| PDGF-BB | B1H0W5 | SLGSPTVAAEPAVIAECKTRTEVFEISRRLIDRTNANFLVWPPCVEVQRCSGCCNNRNVQCRPTQVQDRKVQVKKIEIVRKKKIFKKATVTLVDHLACRCETVVA | *Bos taurus* (Bovine) |
| PDGF-BB | R4GCL3 | SLDGEEPVETAVLAECKTRIEVFEITRSMVDSTNANFLVWPPCVEVQRCTGCCNTRSMQCRPTQARVRHVQVNKIEIVLRKPVFHKAVVALEDHLECRCEPVSS | *Anolis carolinensis* (Green anole) |
| PDGF-BB | A0A3N0Y9E6 | AQPAQQAICKIRTEVIEVTRSMLDRSNANFLLWPPCVEVQRCSGCCNTKTLQCVPVLTHTRYLQVMKIQYVNKRPLYDKAVVSVLDHVECRCQPAP | *Anabarilius grahami* (Kanglang fish) |
| PDGF-BB | A0A3Q1HI22 | AQPAQQAACKVRTEVMEVTRSMLDRRNADFMLWPPCVEVQRCSGCCNTRQLKCVPTVTSKRYLQVTKIQFINRKPHYEKAIISVEDHVSCRCQPAS | *Anabus testudineus* (climbing perch) |
| PDGF-BB | A0A556TV19 | AQPAQQAQCKVRTEVMEVTRSMLDRSNANFLLWPPCVEVQRCSGCCNTKNLKCVAVLTHTRYLQVMKIEYVNMRPVYNKAVVSVNDHVECRCQPAP | *Bagarius yarrelli* (Devil catfish) |
| PDGF-BB | XP_042173650.1 | AQPAQQAVCKVRTEVMEVTRAMLDRRNANFLLWPPCVEVQRCSGCCNTRMLQCVPTVTQTRYLQVTRIQYIDKRPHYDKAVISVEDHASCRCQTHP | *Oncorhynchus tshawytcha* (Chinook salmon) |

| Growth factor | GenBank Acc. No. | Amino acid sequence | Organism |
|---|---|---|---|
| IGF-1 | Q14WA7 | GPETLCGAELVDALQFVCGDRGFYFSKPTGYGSSSRRLHHKGIVDECCFQSCDLRRLEMYCAPIKPPKSA | *Anser anser* (domestic goose) |

| | | | |
|---|---|---|---|
| IGF-1 | A0A1L8GUV7 | GPETLCGAELVDTLQFVCGDRGFYFSKPTGYGSNNRRSH<br>HRGIVDECCFQSCDFRRLEMYCAPAKQAKSA | *Xenopus laevis* (African clawed frog) |
| IGF-1 | P07455 | GPETLCGAELVDALQFVCGDRGFYFNKPTGYGSSSRRAP<br>QTGIVDECCFRSCDLRRLEMYCAPLKPAKSA | *Bos taurus* (Bovine) |
| IGF-1 | BAF74503.1 | GTETLCGAELVDTLQFVCGDRGFYFSKPTGYGSSSRRSH<br>NRGIVDECCFQSCELRRLEMYCAPVKPGKAA | *Anguilla japonica* (Japenese eel) |
| IGF-1 | AEA36762.1 | GPETLCGAELVDTLQFVCGERGFYFSKPTGYGPNARRPH<br>NRGIVDECCFQICELRRLEMYCAPAKTSKAA | *Gadus morhua* (*Cod* ) |
| IGF-1 | Q02815 | GPETLCGAELVDTLQFVCGERGFYFSKPTGYGPSSRRSHN<br>RGIVDECCFQSCELRRLEMYCAPVKSGKAA | *Oncorhynchus tshawytcha* (Chinook salmon) |
| IGF-2 | P01344 | AYRPSETLCGGELVDTLQFVCGDRGFYFSRPASRVSRRSR<br>GIVEECCFRSCDLALLETYCATPAKSE | *Homo sapiens* (Human) |
| IGF-2 | B8QGI3 | AYRPSETLCGGELVDTLQFVCGDRGFYFSRPSSRINRRSR<br>GIVEECCFRSCDLALLETYCATPAKSE | *Bos taurus* (Bovine) |
| IGF-2 | B0Z6G5 | AETLCGGELVDALQFVCEDRGFYFSRPTSRGNNRRPQTR<br>GIVEECCFRSCDLNLLEQYCAKPAKSE | *Oreochromis niloticus* (Nile tilapia) |
| TGFb-1 | P01137 | ALDTNYCFSSTEKNCCVRQLYIDFRKDLGWKWIHEPKGY<br>HANFCLGPCPYIWSLDTQYSKVLALYNQHNPGASAAPCC<br>VPQALEPLPIVYYVGRKPKVEQLSNMIVRSCKCS | *Homo sapiens* (Human) |
| TGFb-1 | P18341 | ALDTNYCFSSTEKNCCVRQLYIDFRKDLGWKWIHEPKGY<br>HANFCLGPCPYIWSLDTQYSKVLALYNQHNPGASAAPCC<br>VPQALEPLPIVYYVGRKPKVEQLSNMIVRSCKCS | *Bos taurus* (Bovine) |
| TGFb-1 | XP_024253659.1 | TTTEEICSDKSESCCVRRLYIDFRKDLGWKWIHEPTGYFA<br>NYCIGPCTYIWNTENKYSQVLALYKHHNPGASAQPCCVP<br>QVLEPLPIIYYVGRQHKVEQLSNMIVKSCRCS | *Oncorhynchus tshawytcha* (Chinook salmon) |
| TGFb-1 | XP_012727903.1 | TSGPETCTAQTENCCVRSLYIDFRKDLGWKWIHKPTGYH<br>ANYCMGSCTYIWNAENKYSQILALYKHHNPGASAQPCC<br>VPQTLDPLPILYYVGRQHRVDQLSNMVVKSCKCS | *Fundulus heteroclitus* (Mummichog) |
| TGFb-1 | P09531 | DLDTDYCFGPGTDEKNCCVRPLYIDFRKDLQWKWIHEPK<br>GYMANFCMGPCPYIWSADTQYTKVLALYNQHNPGASA<br>APCCVPQTLDPLPIIYYVGRNVRVEQLSNMVVRACKCS | *Gallus gallus* (Chicken) |
| TGFb-1 | A0A091JRM2 | RKKRALDAAYCFRNVQDNCCLRPLYIDFKRDLGWKWIHE<br>PKGYHANFCAGACPYLWSSDTQHSRVLSLYNTINPEASA<br>SPCCVSQDLEPLTILYYIGKTPKIEQLSNMIVKSCKCS | *Egretta garzetta* (Little egret) |
| TGFb-1 | XP_029026519.1 | DTRDTCTAQTDSCCVRSLYIDFRKDLGWKWINKPTGYH<br>ANYCMGSCTYIWNAENKYSQILALYKHHNPGASAQPCC<br>VPQTLEPLPILYFVGRQHKVEQLSDMIVKSCKCS | *Betta splendens* (Fighting fish) |
| TGFb-1 | QDA39851.1 | ALDAAFCSRNVQDNCCLRSLYIDFKKDLGWKWIHEPKGY<br>NANFCAGACPYLWSADTQHSNILGLYNTINPEASASPCC<br>VSQDLEPLTILYYIGKNPKIEQLSNMIVKSCKCS | *Carassius gibelo* (Prussian carp) |
| TGFb-1 | AAN03842.1 | TETKDTCTAQTETCCVRSLYIDFRKDLGWKWIHKPTRYH<br>ANYCMGSCTYIWNAENKYSQILALYKHHNPGASAQPCC<br>VPQALEPLPILYYVGRQHKVEQLSNMIVKSCKCS | *Sparus aurata* (Gilthead bream) |

| Growth factor | GenBank Acc. No. | Amino acid sequence | Organism |
|---|---|---|---|
| GDF-8 (MSTN) | O14793 | DFGLDCDEHSTESRCCRYPLTVDFEAFGWDWIIAPKRYK<br>ANYCSGECEFVFLQKYPHTHLVHQANPRGSAGPCCTPTK<br>MSPINMLYFNGKEQIIYGKIPAMVVDRCGCS | *Homo sapiens* (Human) |
| GDF-8 (MSTN) | O18836 | DFGLDCDEHSTESRCCRYPLTVDFEAFGWDWIIAPKRYK<br>ANYCSGECEFVFLQKYPHTHLVHQANPRGSAGPCCTPTK<br>MSPINMLYFNGEGQIIYGKIPAMVVDRCGCS | *Bos taurus* (Bovine) |
| LIF | P15018 | SPLPITPVNATCAIRHPCHGNLMNQIKNQLAQLNGSANAL<br>FISYYTAQGEPFPNNVEKLCAPNMTDFPSFHGNGTEKTKL<br>VELYRMVAYLSASLTNITRDQKVLNPTAVSLQVKLNATID<br>VMRGLLSNVLCRLCNKYRVGHVDVPPVPDHSDKEAFQRK<br>KLGCQLLGTYKQVISVVVQAF | *Homo sapiens* (Human) |
| LIF | Q27956 | SPLPITPVNATCATRHPCPSNLMNQIRNQLGQLNSSANSLF<br>ILYYTAQGEPFPNNLDKLCSPNVTDFPPFHANGTEKARLV<br>ELYRIIAYLGASLGNITRDQKVLNPYAHGLHSKLSTTADVL<br>RGLLSNVLCRLCSKYHVSHVDVTYGPDTSGKDVFQKKKL<br>GCQLLGKYKQVIAVLAQAF | *Bos taurus* (Bovine) |
| LIF | P09056 | SPLPITPVNATCAIRHPCHNNLMNQIRSQLAQLNGSANAL<br>FILYYTAQGEPFPNNLDKLCGPNVTDFPPFHANGTEKAKL<br>VELYRIVVYLGTSLGNITRDQKILNPSALSLHSKLNATADIL<br>RGLLSNVLCRLCSKYHVGHVDVTYGPDTSGKDVFQKKKL<br>GCQLLGKYKQIIAVLAQAF | *Mus musculus* (Mouse) |
| EGF | XP_029020455.1 | NSVESCPATHQSYCLYQGICFYFPEMDSYACTCLPGYIGE<br>RCQFSDLEWWELQ | *Betta splendens* (Fighting fish) |

**Supplementary Table 3.** Cost analysis breakdown (COGS) for recombinant growth factor production on a "laboratory scale". *Capital costs of essential laboratory equipment (e.g., floor shakers, centrifuge, sonicator, benchtop centrifuge, autoclave, dishwashing, gel casting system, electricity etc.) not included.*

| Consumable description | Quantity used | Cost (CAD) | Unit specification | Total cost (CAD) |
|---|---|---|---|---|
| **PROTEIN PURIFICATION CONSUMABLES COSTS** | | | | |
| **(12 L of protein grown by "shaker-flask approach"; 4L erlenmeyer flasks)** | | | | |
| *(All cost above reflected for 12L culture purification)* | | | | |
| Ni-NTA resin superflow (Qiagen) NEW | 3 mL total/target purified | $ 450.00 | 25 mL | $ 90.00 |
| HEPES (Bioshop) | 11.9 g/ L for 50 mM | $ 270.00 | 500 g | $ 6.43 |
| Sodium chloride (Bioshop) | 17.6 g/ L for 0.3 M | $ 50.00 | 10 kg | $ 0.09 |
| Glycerol (Bioshop) | 50 mL | $ 25.00 | 1000 mL | $ 1.25 |
| Imidazole (Bioshop) | 0.35 g/ L for 5 mM | $ 82.00 | 500 g | $ 0.06 |
| (cost for making **1L buffers** for affinity purification) | 2 g/ L for 30 mM | | | $ 0.33 |
| | 17 g/ L for 250 mM | | | $ 2.79 |
| TB broth (Terrific broth culture medium) | 48 g powder for 1 L | $ 60.00 | 500 g | $ 69.12 |
| Vivaspin concentrator (GE Biosciences) | 1 no./ per target | $ 180.00 | 12 nos. | $ 15.00 |
| 50 mL Falcon tubes (VWR) | 15 nos. | $ 110.00 | 500 nos | $ 3.30 |
| Snakeskin dialysis (30.5 cm=1 foot); ThermoFisher | 30 cm | $ 235.00 | 35 feet (35 x 30.5 cm) | $ 6.60 |
| Bradford reagent (BioRad) | 5 mL | $ 190.00 | 500 mL | $ 1.90 |
| Chromatography gravity columns (BioRad) | 2 no. | $ 225.00 | 4 nos. | $ 112.50 |
| Acrylamide (Bioshop) | 4 mL/gel cast | $ 56.00 | 500 mL | $ 0.45 |
| TEMED (Bioshop) | 0.02 mL/gel cast | $ 38.00 | 50 mL | $ 0.02 |
| APS (Bioshop) | 0.5 mL (10%)/gel cast | $ 14.00 | 25 g | $ 0.56 |
| Disodium hydrogen phosphate (Sigma) | 2.88 g/2 L buffer (x2 for 4L buffer) | $ 110.00 | 500 g | $ 1.27 |
| Potassium dihydrogen phosphate (EMD) | 0.5 g/2 L buffer | $ 970.80 | 500 g | $ 1.94 |
| Sodium chlordie (Bioshop) | 16 g/2 L buffer | $ 50.00 | 10 kg | $ 0.16 |
| Potassium chloride (Bioshop) | 0.4 g/2 L buffer | $ 91.00 | 500 g | $ 0.15 |
| **(PBS buffer usage per 2 x 2L)** | | | | |
| **Labour costs** | **24h** approx. (8h x 3 days) | $ 38.00 | per hour | $ 912.00 |
| | **TOTAL COST (for 12L scale up purification; 120 mg total protein yield )** | | | **$ 1,225.90** |
| If protein yields | were **10 mg protein/ L**; so for 12L, **10 mg x 12 litre = 120 mg** | | | |

**CLONING CONSUMABLES COSTS (estimated for 96 targets, high throughput set up)**

| Consumables description | Quantity used | Cost (CAD) | Unit specification | Total cost for 96 targets (CAD) | Total cost/target (CAD) |
|---|---|---|---|---|---|
| * 440 bp for FGF-2; 340bp for TGFB1, 320 bp for PDGF-BB | | | | | |
| Gene synthesis/96 targets* (Twist BioSciences) | **400 bp** approx. bases/protein target | $ 0.08 | base | $ 3,225.60 | $ 33.60 |
| Sequencing primers forward/96 well plate (EuroFins) | 0.10/base | $ 300.00 | 0.10/base | $ 300.00 | $ 3.13 |
| Sequencing primers reverse/96 well plate (EuroFins) | 0.10/base | $ 300.00 | 0.10/base | $ 300.00 | $ 3.13 |
| TCAG sequencing facility/target | $3.50 x 2 reactions | $ 3.50 | per reaction | $ 672.00 | $ 7.00 |
| Miniprep 96-well plate (Qiagen) | | $ 300.00 | per 96 well plate | $ 300.00 | $ 3.13 |
| PCR purification 96-well plate (Qiagen) | | $ 200.00 | per 96 well plate | $ 200.00 | $ 2.08 |
| Miscellaneous: cloning buffers, Pfx, Taq polymerase, dNTP mix, sspI enzyme, T4 polymerase) | | $ 100.00 | | $ 100.00 | $ 1.04 |
| **Labour costs** | **16h** approx. (8h x 2 days) | $ 38.00 | per hour | $ 608.00 | $ 6.33 |
| | | | | **$ 5,097.60** | **$ 59.43** |
